# A knowledge–based scoring function to assess the stability of quaternary protein assemblies

**DOI:** 10.1101/562520

**Authors:** Abhilesh S. Dhawanjewar, Ankit A. Roy, M.S. Madhusudhan

**Author notes:** Equal contribution (Abhilesh S. Dhawanjewar, Ankit A. Roy).

## Abstract

**Motivation:** The elucidation of all inter protein interactions would significantly enhance our knowledge of cellular processes at a molecular level. Given the enormity of the problem, the expenses and limitations of experimental methods, it is imperative that this problem is tackled computationally. *In silico* predictions of protein interactions entail sampling different conformations of the purported complex and then scoring these to assess for interaction viability. In this study we have devised a new scheme for scoring protein-protein interactions.

**Results:** Our method, PIZSA (Protein Interaction Z Score Assessment) is a binary classification scheme for identification of stable protein quaternary assemblies (binders/non-binders) based on statistical potentials. The scoring scheme incorporates residue-residue contact preference on the interface with per residue-pair atomic contributions and accounts for clashes. PIZSA can accurately discriminate between native and non-native structural conformations from protein docking experiments and outperform other recently published scoring functions, demonstrated through testing on a benchmark set and the CAPRI Score_set. Though not explicitly trained for this purpose, PIZSA potentials can identify interactions that are crystallization artefacts.

**Availability:** PIZSA is implemented as a webserver at http://cospi.iiserpune.ac.in/pizsa/.

**Supplementary information:** Supplementary data are available at *Bioinformatics* online.

## 1 Introduction

Proteins, often referred to as ‘biology’s workforce’, rarely act in isolation. More than 80% of all proteins in a cell interact with one another and with other biomolecules to bring about cellular functions such as mediating signal transduction, translating energy to physical motion, immunological response, enzymatic reactions and a host of other cellular processes (Berggård et al., 2007). Proteins function in a crowded cellular environment, diffusing randomly and colliding with one another. Only a small fraction of these collisions result in biologically meaningful associations. Disrupting or interfering with these associations or complexes could lead to disease conditions such as cancer or neurodegenerative conditions in humans (Ryan and Matthews, 2005; Kuzmanov and Emili, 2013). Unsurprisingly, identification and characterization of protein-protein interactions is a fundamental problem in biology that is a necessary step for a complete understanding of the full repertoire of cellular pathways (Vazquez et al., 2003; Stelzl et al., 2005; Zhang et al., 2012).

A variety of experimental methods, based on either biophysical, biochemical or genetic principles have been developed to detect protein-protein interactions (PPIs) (see Zhou et al., 2016 for a comprehensive review). These methods however, are expensive, labor-intensive, and often have additional limitations. Hence, there is considerable interest towards the development of computational approaches to predict and analyze protein-protein interactions (Aloy and Russell, 2006). The computational predictions usually have to surmount two challenges-sampling and scoring. While different computational approaches are utilized in constructing plausible models (sampling) of interacting proteins (see Soni and Madhusudhan, 2017 for a review of methods), a crucial aspect of these predictions is to correctly recognize (scoring) the native (or near-native) interaction from among the models sampled. Ideally, the scoring scheme would rank order the predicted conformations to match what is or should be experimentally observed. Scoring schemes are usually based on free energy calculations incorporating different atomic interactions (Pierce and Weng, 2007; Lyskov and Gray, 2008) or on statistical methods (also referred to as statistical potentials) that extract information from known protein interactions (Huang and Zou, 2008; Rykunov and Fiser, 2010; Liu and Vakser, 2011). Some scoring scheme even combine these two methods (Zimmermann et al., 2012). In addition to this there are machine learning techniques to score/evaluate possible interactions conformations (Bordner and Gorin, 2007).

The premise of the statistical potentials is that the ratio of an observation (of interaction) to its expectation is indicative of interaction strength. The higher the ratio, the greater is the binding energy. The key to these computations is in making a precise estimate of the expectation value. The initial methods were simple and attempted to convert the observed/expected ratios into free energies (Tanaka and Scheraga, 1976; Sippl, 1995). These potentials got increasingly more nuanced with the inclusion of additional considerations such as solvent effects and Lennard-Jones potentials (Miyazawa and Jernigan, 1996; Tovchigrechko and Vakser, 2006).

In study we have devised a new statistical potential. Here, we derive amino acid interaction preference matrices from three-dimensional protein structures in the Protein Data Bank (Berman et al., 2000) that extends the concept employed in Davis et al., 2006. We use these preference matrices to propose a new scoring function for the binary classification of protein assembly stability as well as for rank-ordering different docking poses for a given protein complex. We refer to the binary classification algorithm as PIZSA for Protein Interaction Z-Score Assessment. Our potential function explicitly includes local geometry and interface propensities while also incorporating the strength of individual residue pair interactions and accounting for clashes. The expected probability function in our formulation also accounts for the relative abundance of different amino acid residues leading to a more appropriate reference state. We compare our scoring function to the CIPS potential in their abilities to discriminate between native and near-native protein complex structures from the Dockground Decoy Set (Kundrotas et al., 2018) CAPRI Score_set (Lensink and Wodak, 2014) and report an improvement in discriminatory performance. Though our potential functions have not been explicitly trained for this purpose, they can also discriminate between biologically meaningful interfaces and crystal artefacts.

## 2 Materials and Methods

### 2.1 Datasets used for construction of statistical potentials

#### 2.1.1 Construction of residue pairing preference matrices

Protein-protein interface residue pairing preference matrices were constructed from the three-dimensional structural data of dimeric protein complexes retrieved from the Protein Data Bank (PDB) (Berman et al., 2000). The dataset was culled using the PISCES (Wang and Dunbrack Jr, 2003) web server to eliminate redundancy and retain structures with a maximum sequence identity of 40%, a minimum resolution of 3 Å and a maximum R-factor of 0.3. After culling, the dataset reduced from 32,871 to a set of 3,497 dimeric complexes (Supplementary S1).

#### 2.1.2 Binary classification

Protein associations are classified as stable or unstable by calculating Z scores. The background distribution for the calculation of Z scores was estimated using 1000 decoy structures each for 351 native protein complexes from the Dockground Docking Decoy Set 2 (Kundrotas et al., 2018). The classification performance was tested on Dockground Docking Decoy Set 1 (Kundrotas et al., 2018), comprising of 61 native complexes with 100 decoys for each native complex.

#### 2.1.3 Ranking native complex

The ability to rank native complexes amongst the best scoring interactions was benchmarked using the CAPRI Score_Set (Lensink and Wodak, 2014) and the Dockground Docking Decoy Set 2. The CAPRI Score_Set consists of 13 dimeric and 2 trimeric target complexes. 322 of 351 targets in Dockground Docking Decoy Set 2 were used for benchmarking as 26 targets had decoys with non-canonical atom names and 3 targets had decoys that we were unable to score using CIPS (Supplementary S2).

#### 2.1.4 Identification of crystal artefacts

We tested the ability of our potentials to distinguish between biologically meaningful interactions and crystal packing artefacts on two datasets. The ‘Bahadur’ (Bahadur et al., 2004) and ‘Duarte’ (Duarte et al., 2012) datasets comprised of 57 and 35 structures of validated crystal contact artefacts respectively (Supplementary S3).

### 2.2 Construction of the composite scoring functions

#### 2.2.1 Scoring function

Scoring matrices were constructed from the ratio of observed probabilities of interface residue pairs to their expected probabilities of occurrence at the interface. Two residues from different protein chains are identified as an interface residue pair if one or more atoms from one residue is within a threshold distance from atoms of the other residue. The score 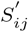 for a residue pair *ij* is calculated as:

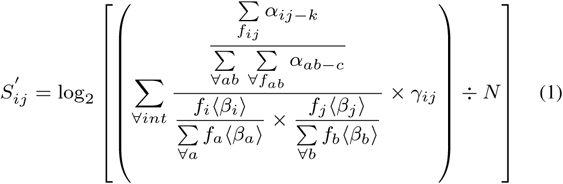

where, *α*_*ij−k*_ is the atomic propensity of interface residue pair *ij* with *k* atoms. All frequencies are observed frequencies. *f*_*ij*_ is the frequency of interface residue pair *ij. ab* is any interface residue pair and *c* is the number of atoms involved in any interface residue pair interaction. ⟨*β*_*i*_⟩ and ⟨ *β*_*j*_ ⟩ are the average atomic propensities of interface residues *i* and *j* respectively. *f*_*i*_ and *f*_*j*_ are the frequencies of interface residues *i* and *j* respectively. *γ*_*ij*_ is the abundance normalization term for residue pair *ij. N* is the total number of interfaces and *int* is any interface. The abundance normalization term *γ*_*ij*_ is calculated as:

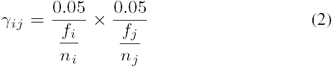

where *n*_*i*_ and *n*_*j*_ are the total number of residues in the monomeric subunits of residues *i* and *j* respectively. The uniform probability of occurrence for any residue is 0.05 (1*/*20).

The atomic propensity, *α*_*ij−k*_ of an interface residue pair *ij* with *k* number of atoms within a threshold distance is calculated as:

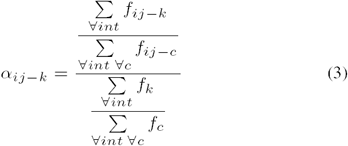

where, *f*_*ij−k*_ is the frequency of residue pair *ij* with *k* number of atoms. *f*_*k*_ is the frequency of any residue pair with *k* number of atoms and *c* is any number of atoms observed in interactions.

Three different scoring matrices were constructed using only main chain atoms, side chain atoms or main chain/side chain atoms exclusive from each residue partner. Favourable interface residue pairs have positive scores whereas unfavorable interface residue pairs have negative scores. Variants of the scoring matrices were constructed at distance thresholds of 4 Å, 6 Å and 8 Å.

#### 2.2.2 Scoring protein-protein complexes

A raw score (*S*_complex_) is assigned to a protein complex by summing up the individual weighted residue pair scores over the interface. Each residue pair receives a score 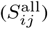 that is a linear combination of the main chain 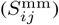, side chain 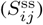 and main chain/side chain 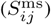 interaction score. Each component of the residue pair score is weighted with an atomic propensity and a clash penalty as:

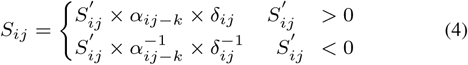

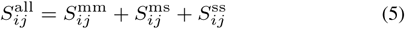

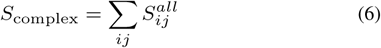

where 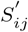 is the unweighted score from any of the three scoring matrices. *δ*_*ij*_ is the clash penalty for residue pair *ij*. 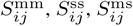 and 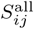 are the weighted main chain, side chain, main chain/side chain and all atom interaction scores respectively. *S*_complex_ is the raw score of the protein complex.

Clash penalty *δ*_*ij*_ is a measure of the severity of atomic clashes in a residue pair interaction. It ranges from 0 for severe clashes to 1 for no clashes and is calculated as:

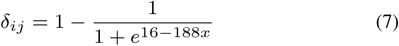

where

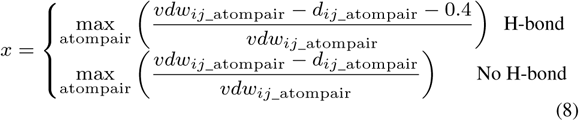

where, *x* is the maximum overlap fraction between interacting atoms that varies from 0 to 1 and is set to 0 when there is no overlap. *vdw*_*ij*_atompair_ is the sum of the van der Waals radii of interacting atom pairs and *d*_*ij*_atompair_ is the Euclidean distance between the interacting atom pairs. Atom pairs that have hydrogen bond donor and acceptor atoms are identified. Such hydrogen bonded atom pairs have an additional tolerance of 0.4 Å. The sigmoidal function used as the clash penalty has been optimized to penalize 20% of the highest overlap fractions observed in the training set such that least number of native complexes are predicted as unstable associations (Supplementary S4).

### 2.3 Binary classification for the stability of protein complexes

The residue preference matrices constructed above were employed to classify protein complexes as stable or unstable associations. To account for the effect of interface size, raw scores for each complex were further normalized by the number of interacting residue pairs (Equation 9). Z score is a measure of how likely a protein complex is to form a stable association. A protein complex is predicted to be a stable binder if the Z score is greater than a threshold. The Z score of protein complexes are calculated as:

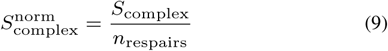

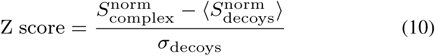

where 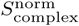 is the normalized raw score for a protein complex. *n*_respairs_ is the number of interacting residue pairs observed in the protein complex. 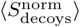 and *σ*_decoys_ are the average and standard deviation of normalized raw scores obtained from background decoys (Dockground Docking Decoy Set 2).

The background distribution for the calculation of Z scores was estimated using 1000 decoy structures each for 336 native protein complexes from the Dockground Docking Decoy Set 2 (Kundrotas et al., 2018). Receiver-Operator Characteristic (ROC) curves were constructed to estimate the observed false-positive and true-positive rates at different Z score thresholds and different distance thresholds. ROCs were then integrated to calculate the area under the curve (AUC) and identify the operating points. Z scores with operating points closest to (0, 1) in the ROC curves were chosen as optimal binary classification thresholds to maximize the True Positive Rate (TPR) and minimize the False Positive Rate (FPR). Classification performance was tested on Dockground Docking Decoy Set 1 and evaluated in terms of accuracy, balanced accuracy and a modified Matthew’s correlation coefficient (MCC) (Equation 11).

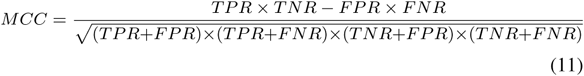

where, TPR is True Positive Rate, FPR is False Positive Rate, TNR is True Negative Rate and FNR is False Negative Rate.

### 2.4 Binding mode selection

The ability of the potential to select the proper binding mode when multiple alternative binding interfaces are present was also assessed. The antigen-binding fragment of camelid antibodies are composed of a single, heavy chain antibody VH domain (VHH). Structures of three different dromedary VHH domains bound to porcine pancreatic *α*-amylase (PPA) at non-verlapping orthogonal sites have been deposited in the PDB (Desmyter et al., 2002). Three-dimernsional structures of non-native binding modes were constructed with homology modeling using MODELLER v9.15 (Šali and Blundell, 1993, Davis et al., 2006) (Supplementary S5). Structures of camelid VHH domains AMB7, AMD9 and AMD10 bound to PPA (PDB codes: 1KXT, 1KXV and 1KZQ respectively) were evaluated for each VHH-PPA complex using the PIZSA potential.

## 3 Results

### 3.1 Construction of the statistical potential matrices

Amino acid pairing propensities capture the likelihood that amino acids *i* and *j* interact across a protein-protein interface. We constructed such propensity matrices defined f or d ifferent a tomic i nteraction categories (main chain-main chain, side chain-side chain and main chain-side chain) at three different distance thresholds (4 Å, 6 Å and 8 Å) (Figure 1). The number of favorable amino acid pairs at the interface were highest for the side chain-side chain mode of interaction followed by main chain-side chain and main chain-main chain interaction. For example, at 4 Å 30% (63 out of 210) of the side chain-side chain, 1.5% (6 out of 400) of the main chain-side chain and 0.5% (1 out of 210) of the main chain-main chain amino acid pairs were favorable. The number of favorable amino acid pairs increases with an increase in distance cut-off. For example, with an increase in distance cut-off from 4 Å to 6 Å and 8 Å the number of favorable side chain-side chain pairs increases from 30% to 57.1% and 71.9%; main chain-side chain increases from 1.5% to 21.3% and 49.3%; main chain-main chain increases from 0.5% to 1.4% and 32.8% respectively. This trend is also observed to be monotonic across all three modes of interactions. At all distance cut-offs, side chain-side chain mode of interaction contributed the most, since most interactions on the interface were side chain mediated interactions. For example, at 4 Å distance cut-off 58% of the interactions were of the side chain-side chain type whereas the main chain-side chain and main chain-main chain account for 34% and 8% of the interactions respectively.

**Fig. 1.**
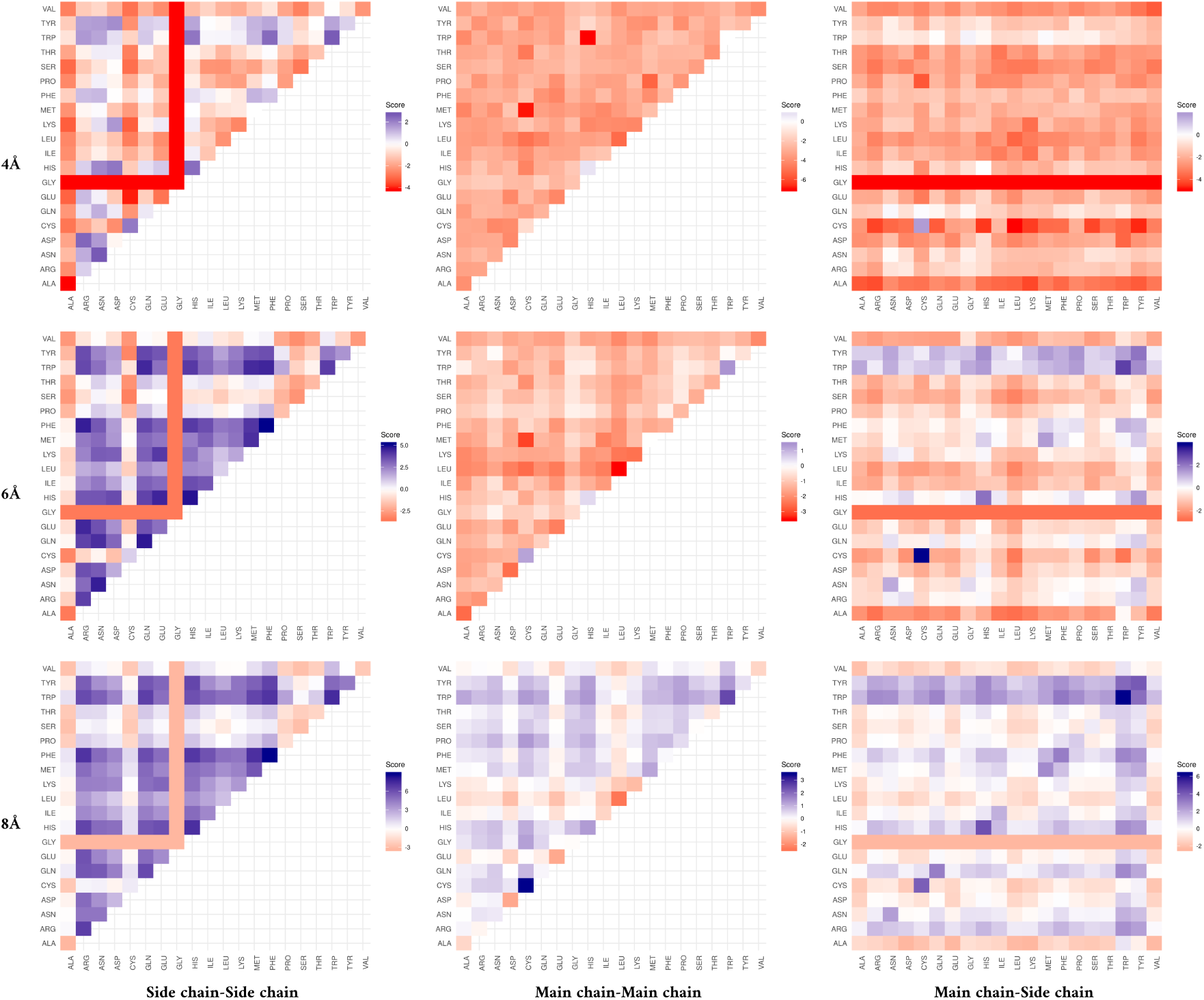
Amino acid propensity matrices constructed at three different interaction distance thresholds (4 Å, 6 Å, and 8 Å) and three different atomic interaction types (side chain-side chain, main chain-main chain and main chain-side chain). The side chain-side chain and main chain-main chain matrices are symmetric whereas the main chain-side chain matrix is asymmetric since the residues exclusively interact either via main chain atoms (x-axis) or side chain atoms (y-axis). Bluer shades represent more favorable interactions whereas redder shades represent less favorable interactions.

Since the residue pairing preferences across different modes of interaction were not identical we identified the contribution from all different modes separately. Further, we estimated the distribution of the number of atomic contacts shared by each residue-residue interaction on the interface and found that many residue pairs have a strong tendency to interact with a preferred number of atomic contacts. Our propensity scores account for this optimal number of atomic contacts (*α*_*ij−k*_, Equation 3). The atomic propensities (*α*_*ij−k*_) are an observed by expected ratio of the probability that an interaction between residues *i* and *j* is mediated by *k* number of atomic contacts. The expected probability that any residue pair interacts through *k* number of atoms declines exponentially with increasing number of atoms. The observed probability distribution fits the expected probability distribution closely for some residue pairs but for other residue pairs the observed probabilities are much higher than their expected probabilities for a certain number of interacting atoms. For example, the observed and expected probability distributions match closely for the ASP-PHE residue pairs whereas ARG-GLU pairs tend to have 6 to 8 atomic contacts (6 being the highest) at 4 Å distance cut-off and side chain-side chain mode of interaction. Interactions with atomic propensities above 1 (observed probabilities higher than the expected) indicate favorable number of atomic contacts whereas those below 1 (observed probabilities lower than the expected) indicate sub-optimal number of atomic contacts (Figure 2, Supplementary S6). ARG-GLU amino acid pairs have higher observed probabilities for 6 to 8 atomic interactions as compared to what is expected (Figure 2). Also ARG-GLU pairs with 2 to 4 atomic interactions are sub-optimal as indicated by the lower observed probability as compared to the expected (Figure 2).

**Fig. 2.**
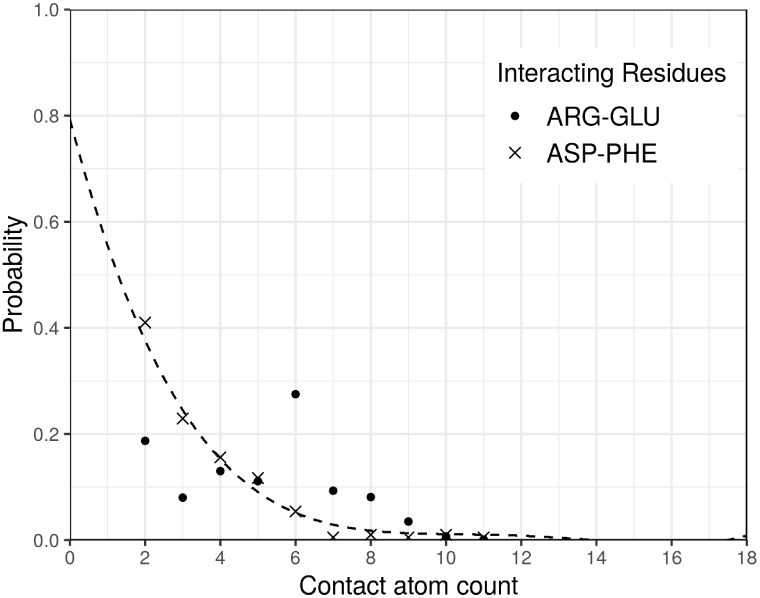
The observed (•, ×) and expected (dashed line) probability distribution profile of the number of atoms mediating residue pair interactions in ASP-PHE and ARG-GLU.

**Fig. 3.**
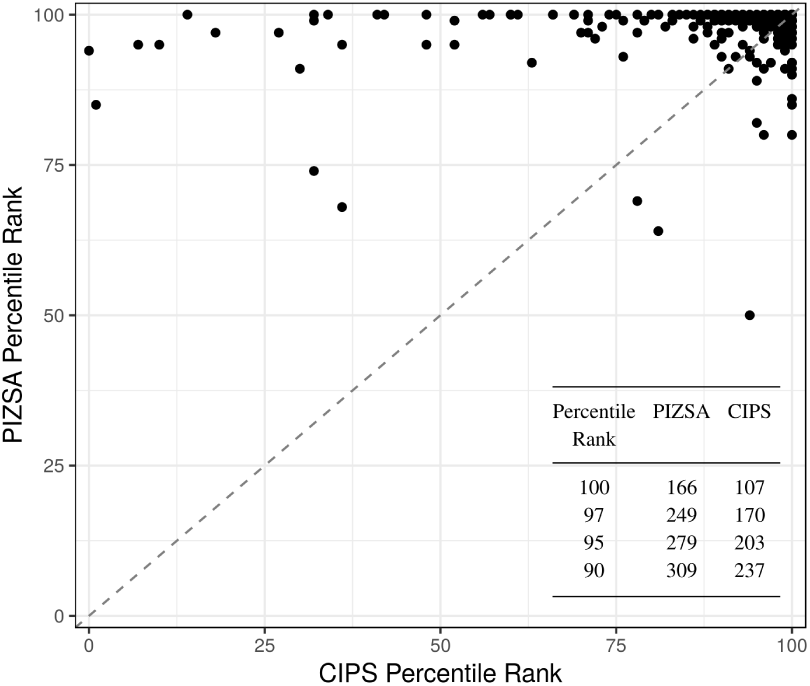
The percentile ranks assigned by PIZSA and CIPS to native structures from the Dockground Docking Decoy Set. Points above/below the diagonal indicate cases where PIZSA assigned a better/worse rank than CIPS. Inset table shows the number of native complexes ranked above different percentiles.

The amino acid pairing preferences exhibited by the PIZSA scoring matrices are qualitatively similar across the three distance thresholds and suggest that: (i) oppositely charged residues have a strong tendency to pair across the interface with a large overlap of atomic contacts, (ii) residues with aromatic rings (HIS, TRP and TYR) have favorable interactions with most amino acids indicating they play a crucial role on interfaces, (iii) interactions among hydrophobic residues were less favorable at 4 Å but more favorable at 6 Å and 8 Å, especially for the side chain-side chain potentials. The side chain-side chain propensity matrices show the highest degree of specificity, followed by the main chain-side chain matrices and then the main chain-main chain matrices across the three distance thresholds. Top three highest scoring residue pairs in the main chain-main chain and main chain-side chain matrices were HIS-HIS, TRP-TRP and CYS-CYS in no particular order. In the side chain-side chain matrices, the top three scoring amino acid pairs included ASN-ASN, TRP-TRP and ARG-ASP at 4 Å, PHE-PHE, HIS-HIS and GLN-GLN at 6 Å and PHE-PHE, TRP-TRP and HIS-HIS at 8 Å. Side chain matrices accounted for many high scoring residue pairs with diverse interaction types. For example, electrostatically interacting residue pairs such as ARG-GLU, hydrogen bonding interactions such as ASN-ASN, *π*-*π* stacking such as TRP-TYR, cation-*π* interactions such as HIS-ARG and hydrophobic interactions such as ILE-ILE. ALA and VAL had the least number of favorable interactions in all cases. Smaller amino acids such as SER, THR, PRO had less favorable interactions except in case of their main chain-main chain interactions at 8 Å whereas larger amino acids such as GLU, ILE, LEU and LYS had less favorable main chain-main chain interactions. CYS-CYS interactions were less favorable in side chain-side chain matrices at 6 Å and 8 Å whereas they were amongst the highest scoring interactions in all other matrices. Side chain-side chain interactions of large aliphatic amino acids LEU and ILE were favorable at 6 Å and 8 Å. Interestingly, ARG-ARG interactions were also favorable in side chain-side chain matrices at higher distance thresholds of 6 Å and 8 Å.

### 3.2 Binary classification of protein complexes

Our scoring matrices describe individual residue pairing preferences on the interface. On an interface containing several interacting residue pairs, we use pair preference values from our scoring matrices in the form of a Z score to identify stable protein-protein associations. Optimum Z score thresholds for classification of protein complexes as stable binders or unstable non-binders were obtained by analyzing receiver operator characteristic (ROC) curves (Supplementary S7). The area under the ROC curves were 0.98, 0.95 and 0.93 for distance thresholds of 4 Å, 6 Å and 8 Å respectively. Optimal Z score thresholds for distance cut-offs of 4 Å, 6 Å and 8 Å are 1.5, 1.2, and 1.0 respectively. The true positive and false positive rates at optimal Z score thresholds are 94.0% and 6.7% for 4 Å, 89.2% and 11.0% for 6 Å, 87.2% and 15.5% for 8 Å respectively. For all further analyses we used 4 Å as the distance cut-off as it had the highest true positive rate and lowest false positive rates of classification.

We tested the classification accuracy of PIZSA on the Dockground Docking Decoy Set 1, which comprises of 61 protein complexes with 100 decoy conformation for every native conformation. As this testing set is skewed with respect to the ratio of true positives to true negatives (1:100). we evaluated the performance by estimating the accuracy, balanced accuracy and a modified Matthew’s correlation coefficient as described in the methods section. PIZSA was able to correctly predict 55 out of 61 natives as stable binders with an accuracy of 0.85, a balanced accuracy of 0.87 and an MCC of 0.75 (Supplementary S8).

Each protein quaternary complex is assigned a score that is the sum of individual scores for residue pair interactions across the protein-protein interface. Each residue pair interaction score is in turn a sum of the residue pair preferences from the three different matrices defined for different atomic interactions (main chain-main chain, main chain-side chain and side chain-side chain) and distance thresholds. Residue pair preference values are further weighted by their atomic propensities (*α*_*ij−k*_) and a clash penalty (*γ*_*ij*_). The clash penalty term is used to assess the severity of atomic clashes in residue pair interactions and proportionally decrease their scores. The inclusion of these terms in the scoring function further refines the ability of our potential to identify native interactions as exemplified here with the insect anti-freeze protein complex (PDB ID: 4DT5). This example illustrates how the inclusion of weighting terms significantly improves the discriminatory performance compared to a standard pair potential. At a distance threshold of 4 Å, the homodimeric complex is exclusively composed of 23 threonine-threonine (THR-THR) residue pairs. THR-THR pair has a main chain-main chain, main chain-side chain and side chain-side chain score of −3.89, −2.98 and −1.30 respectively. In the absence of the weighting terms, each THR-THR pair would receive a score of −8.17. The cumulative score for the interface would then be −187.91 with a Z score of −0.44 and the interface would be classified as a non-binder. However, the inclusion of the weights increase the cumulative score to −30.34, the Z score to 2.78 and predicting the interface as a binder.

### 3.3 Identification of native protein complexes and comparison with CIPS potentials

Recently, a new amino acid pairwise interaction potential, CIPS (Nadalin and Carbone, 2018), was proposed to rank-order different docking configurations. It takes into account a contact based measure that weighs residue pairing frequencies by the number of atomic contacts shared between the residue pair. When compared to three previously published residue preference matrices (Glaser et al., 2001; Pons et al., 2011; Mezei, 2015), CIPS was found to be better at discriminating high-quality structural models from decoys than others.

In this article, we compare ourselves with CIPS as they have been shown to outperform three other statistical potentials in rank-ordering protein-protein interaction complexes. Our method differs from that of CIPS in five essential ways: (i) we have constructed three mutually exclusive amino acids preference matrices that categorize interactions according to the type of atoms involved in an interaction; (ii) we do not make use of explicit solvent accessibility calculation and the amino acids preferences are solely based on the distribution of residues and their interacting atoms in Euclidean space; (iii) we account for atomic propensities similar to CIPS’ contact propensity but with a completely different reference state; (iv) we introduced a penalty for steric clashes that scales scores according to the severity of the clash, and lastly (v) we have introduced a measure that classifies a protein-protein complex as a stable or unstable association.

#### 3.3.1 Performance on the Dockground decoy set

One of the objectives of protein docking experiments is to identify the native interactions from a set of decoys. This is usually achieved with the help of a scoring function that utilises features of interfaces such that native complexes score optimally. One class of such scoring functions, employed by both PIZSA and CIPS, make use of amino acid contact propensities derived from known structures of protein complexes. Previously, CIPS had compared the performance of their scoring scheme to other scoring schemes (Glaser et al., 2001; Pons et al., 2011; Mezei, 2015) on the Dockground Docking Decoy Sets and CAPRI decoy sets. Here, we compare the performance of PIZSA to CIPS on Dockground Docking Decoy Set 2. The ability of PIZSA pair potentials to discriminate the native structural conformation was compared with CIPS potentials on a testing set comprising of 322 native structures with 100 decoy structures each from the Dockground Decoy Set 2 (Supplementary S9). PIZSA ranked the native structure as the best (rank 1), in top 3, in top 5 and in top 10 for 166 (52%), 249 (77%), 279 (87%) and 309 (96%) protein complexes respectively whereas CIPS ranked the native structure as the best (rank 1), in top 3, in top 5 and in top 10 for 107 (33%), 170 (53%), 203 (63%) and 237 (74%) protein complexes respectively (Table 1). Further for individual comparisons, PIZSA ranked 175 structures better than CIPS, 75 structures equal to CIPS and 72 structures worse than CIPS. For 65 of the 72 structures where the CIPS rank is better, PIZSA scores the native structure not more than 10 ranks below CIPS rank.

**Table 1.** Rank-ordering of the native complexes for targets in the CAPRI Score_set

| Target | PDB ID | Number of decoys | PIZSA rank<sup>a</sup> | CIPS rank<sup>b</sup> |
| --- | --- | --- | --- | --- |
| T29 | 2VDU | 2016 | 76.88 | 93.33 |
| T30 | 2REX | 1119 | 94.82 | 96.13 |
| T32 | 3BX1 | 599 | 99.50 | 45.74 |
| T35 | 2W5F | 499 | 100.00 | 23.25 |
| T36 | 2W5F | 309 | 100.00 | 3.24 |
| T37 | 2W83 | 1495 | 99.60 | 76.86 |
| T38 | 3FM8 | 888 | 98.31 | 80.47 |
| T39 | 3FM8 | 1386 | 99.64 | 82.14 |
| T40 | 3E8L | 2144 | 75.93 | 61.42 |
| T41 | 2WPT | 1180 | 72.03 | 28.83 |
| T46 | 3Q87 | 1640 | 78.54 | 85.17 |
| T47 | 3U43 | 1051 | 84.49 | 96.48 |
| T50 | 3R2X | 1448 | 98.76 | 89.94 |
| T53 | 4JW2 | 1400 | 78.43 | 94.71 |
| T54 | 4JW3 | 1398 | 74.32 | 48.79 |

<sup>a, b</sup> % decoys that score worse than the native.

#### 3.3.2 Performance on the CAPRI Score_set

The consolidated benchmark set made available from the CAPRI experiments (CAPRI Score_set) serves as another independent decoy set to benchmark the rank-ordering abilities of scoring functions. The benchmark set contains roughly 19,000 predicted complexes for 15 published CAPRI targets, of which 13 are dimeric complexes and 2 are trimeric complexes (Lensink and Wodak, 2014). Once again we compared our performance in rank ordering the native complexes with that of CIPS.The percentile ranks for the 15 targets assigned by PISZA and CIPS are reported in Table 2. PIZSA assigns better ranks than CIPS for 10 out of 15 targets with 8 native targets ranked in the top 10 percentile and all the native targets ranked in the top 28 percentile. CIPS in comparison assigns top 10 percentile ranks to 4 target structures.

**Table 2.** Z scores of VHH-Porcine Pancreatic *α* amylase complexes

|  | AMB7 mode | AMD10 mode | AMD9 mode |
| --- | --- | --- | --- |
| AMD9 | 0.85 | 1.84 | **2.18** |
| AMD10 | 1.23 | **2.30** | 0.14 |
| AMB7 | **2.23** | 1.73 | 0.78 |

Scores for native complexes are in bold.

We were able to successfully identify 14 out of 15 CAPRI targets (native complexes) as stable associations. Target 40 (3E8L, receptor chain: C, ligand chain: A/B) was predicted as a non-binder with a Z Score of 1.35 which is below the threshold of 1.5. In the 7 cases where PIZSA was unable to rank the native complex in the top 10%, the difference in Z Scores of the native complexes from the best ranked decoys range from 0.27 to 1.34 with 0.76 being the mean difference. Z Score differences between the best decoys and their respective native complexes are less than 1 for T29 (0.94), T41 (0.82), T47 (0.39), T53 (0.27) and T54 (0.50) which implies that the best scoring decoy is within a standard deviation of the native complex score. However, T40 and T46 have Z score differences of 1.34 and 1.10 respectively which implies that the best decoys are more than a standard deviation away from the native complex score. Interestingly, both the targets T40 (3E8L) and T46 (3Q87) have an interaction predominantly mediated by beta sheets. We have also tested our method’s ability to identify near native structures from the CAPRI Score_set and compared our performance with that of CIPS (Supplementary S10 (a) and S10 (b)). Furthermore, PIZSA classified high, medium and acceptable quality near native complexes as stable association with an accuracy of 0.62 on the same dataset (Supplementary S10 (c)).

### 3.4 Binding mode selection

The statistical potentials were also able to select the native binding mode for all three VHH domains (AMB7, AMD9 and AMD10) interacting with porcine pancreatic *α*-amylase (Desmyter et al., 2002). The VHH domains bind to orthogonal sites on the procine pancreatic *α*-amylase despite sharing a high structural similarity with C*α* RMSD ranging from 0.61 to 0.84 Å. We built models for each of the VHH domains interacting with their non-native epitopes on porcine pancreatic *α*-amylase (Davis et al., 2006) and scored them with PIZSA along with their native complexes (Table 2). The native binding modes were successfully identified as the highest scoring interfaces amongst the models. It should be noted that 2 of the 6 non-native binding modes are also scored as viable binders. However, the Z Scores of the native interactions are higher as compared to both of these non-native interactions.

### 3.5 Identification of crystallization artefacts

Distinguishing whether structures of protein assemblies solved by X-ray crystallography are biologically meaningful or simply artefacts of the crystallization process is an important problem in protein-protein interaction studies. Although crystallographic interfaces have smaller surface areas than biologically relevant structures, this is not frequently the case and significant overlap in interface areas has been observed. We tested PIZSA’s ability to distinguish between artefacts and biological structures on two datasets (Bahadur et al., 2004; Duarte et al., 2012) of validated protein crystals with large interface areas. PIZSA was able to correctly identify 45 out of 57 structures (78.94%) and 23 out of 35 structures (65.71%) for the ‘Bahadur dataset’ and the ‘Duarte dataset’ respectively. Upon detailed inspection of cases where PIZSA fails to correctly classify the structures, we found that 6 out of 12 structures from the ‘Bahadur dataset’ and 4 out 12 structures in the ‘Duarte dataset’ had atleast one multimeric assembly in the ‘BIOLOGICAL_UNIT’ record of the PDB entries. Since PIZSA classified these structures as stable multimeric assemblies, these cases represent correct classifications. Including these in the accuracy calculations, PIZSA has a success rate of 89.47% on the ‘Bahadur dataset’ and 77.14% on the ‘Duarte dataset’.

## 4 Discussion

We constructed statistical potentials to assess the stability of protein quaternary assemblies based on amino acid preferences extracted from a large dataset of experimentally deduced protein-protein interfaces. Defining amino acid preferences for different categories of atomic interactions (main chain-main chain, main chain-side chain and side chain-side chain) and at different distance thresholds helped dissect the specific modes through which residues interact across protein interfaces. These residue pairing preferences were further refined by incorporating the relative abundance of amino acid residues in proteins. We observe that residue pairings across the interface tend to occur with a preferred number of shared atomic contacts and incorporate this as an atomic propensity parameter in our scoring function. The inclusion of this parameter also enables us to capture the specificity in atomic interactions between different residue types across the protein-protein interface. We also include a clash penalty to correct for steric clashes in protein structures that can lead to spurious contacts and confound the signal from residue preferences. Our scoring function is able to discriminate stable protein assemblies from unstable assemblies with an accuracy of 0.85, a balanced accuracy of 0.87 and an MCC of 0.75 on the Dockground Docking Decoy Set 1.

The construction of amino acid preference matrices also enable us to identify crucial residues involved in protein binding. Residues containing aromatic rings (TYR, TRP, PHE and HIS) are assigned favorable scores for their interactions with most residues suggesting their versatility in mediating residue pair interactions across the interface. These amino acid residues are characterized by the presence of a *π*-electron cloud above and below the aromatic ring that can interact with other aromatic and non-aromatic residues imparting stability (Ma and Dougherty, 1997; Makwana and Mahalakshmi, 2015). We find that interactions between hydrophobic residues were deemed less favorable at shorter distance thresholds but as more favorable at larger distance thresholds, especially for side chain-side chain interactions of large aliphatic amino acids such as LEU and ILE, an observation that has previously been reported (Bahar and Jernigan, 1997). Electrostatic interactions between oppositely charged residues also had favorable interaction scores and have been found to play an important role in determining specificity in protein interfaces (Sheinerman et al., 2000). These interactions were also frequently found to have high atomic propensities for a specific number of interacting atoms. These patterns are supported by previous observations that the formation of salt bridges between oppositely charged amino acids exhibit well-defined geometric preferences (Donald et al., 2011). For interactions between similarly charged residues, we find that interactions between positively charged residues are more favorable compared to interactions between negatively charged residues which were predominantly unfavorable. Among the positively charged residues, we confirm observations from previous studies that ARG-ARG pairs are more favorable than LYS-LYS pairs (Glaser et al., 2001; Nadalin and Carbone, 2018). Arginine along with tryptophan and tyrosine exhibit strong favorable interactions with most other amino acid types. This could be due to arginine’s capability to form multiple types of favorable interactions, similar to the aromatic residues. In addition to the capability to form H-bonds and salt bridges via its positively-charged guanidinium motif, the electron delocalization of its guanidinium *π*-system gives it a pseudo-aromatic character (Crowley and Golovin, 2005). The presence of three methylene carbon atoms in its side chain also enables arginine to participate in hydrophobic interactions.

The qualitative patterns of amino acid preferences across interfaces extracted with PIZSA potentials differ slightly than those described previously (Keskin et al., 1998; Glaser et al., 2001; Nadalin and Carbone, 2018). However, these patterns are in close agreement with amino acid preferences found in protein interaction hot-spots (Bogan and Thorn, 1998) and hence better represent the specificity in interactions between types of interactions across protein-protein complexes.

Using the Dockground Docking Decoy Set and the CAPRI Score_set, we report a significant improvement for the effective rank-ordering of native conformations over the recently published CIPS potentials, which reported better performance compared to three other propensity matrices and two atomic potentials (Nadalin and Carbone, 2018). Even though the CIPS potential also accounts for residue preferences and the level of connectivity for interface residue pairs, we believe the difference in performance is due to the definitions of interaction strengths in the two models. CIPS defines the interaction strength as the average number of contacts per residue while PIZSA defines the interactions strengths for each interface residue pair based on the ratio of the observed probability to expected probability of number of atomic contacts shared by the residue pair. This definition helps further refine the weights assigned to residue preference scores and in combination with the explicit inclusion of a clash penalty term results in a better discriminatory performance. However, our scoring function was unable to assign optimal scores for two of the CAPRI targets where the interactions predominantly consisted of beta sheets. We believe that PIZSA was unable to assign optimal scores to these interactions since our main chain-main chain scoring matrices contribute minimally to the overall score of an interaction, whereas inter-chain beta sheets form multiple main chain hydrogen bonds that impart stability (Dou et al., 2004).

The ability of PIZSA potentials to distinguish between favorable and unfavorable binding modes is demonstrated through the example of VHH domains in complex with porcine pancreatic *α*-amylase. The three VHH domains have distinct binding modes for complexation with porcine pancreatic *α*-amylase and PIZSA potentials were able to select the native binding modes for all three VHH domains. In addition to effective rank-ordering of near-native structures and the binary classification of protein assemblies, testing on two distinct datasets, PIZSA potentials were also successful in distinguishing between biologically meaningful complexes and crystallization artefacts without explicitly learning from a dataset of structures with crystal artefacts.

We demonstrate that knowledge-based potentials based on known protein-protein interactions capture crucial information about protein binding and can be successfully applied to identify biologically meaningful protein complexes in protein docking experiments, protein-protein interaction predictions and also in distinguishing complexes formed as artefacts of the protein crystallization process. Such potentials could also aid in the design of non-canonical protein complex.

## Acknowledgements

We would like to thank members of the COSPI lab at IISER Pune for valuable discussions and insights.

