## Supplementary Data for "A knowledge–based scoring function to assess the stability of quaternary protein assemblies"

**Supplementary S1.** List of protein complex structures (PDB codes) used for construction of residue pairing preference matrices.

1A25 1A2O 1A6J 1AAZ 1AC6 1ADU 1AMU 1AOE 1AOH 1AOR 1AQU 1AT3 1ATZ 1AU1 1AYO
1AZT 1AZW 1B3U 1B43 1B88 1BC5 1BCM 1BEH 1BF6 1BIN 1BJA 1BMT 1BQU 1BXT 1BYF
1C3R 1C94 1C9O 1CI4 1CI9 1CJA 1CKU 1COL 1COZ 1CP2 1CQ3 1CRU 1CZP 1D0Q 1D2O
1DBW 1DDV 1DEB 1DEK 1DJ7 1DJT 1DLE 1DM9 1DNP 1DOW 1DQE 1DVK 1DWU 1DYN 1DYO
1DYS 1DZK 1E0B 1E30 1E5R 1E6C 1E8C 1E9G 1EAJ 1ECE 1EDM 1EEJ 1EEO 1EGA 1EI7
1EJD 1EJF 1EK6 1EKE 1EM9 1EPA 1EQ9 1EUJ 1EUV 1EXT 1EYV 1EZG 1F08 1F0K 1F0L
1F1C 1F35 1F39 1F46 1F6B 1F7D 1F86 1F9M 1FIW 1FJ2 1FJR 1FM0 1FMT 1FN8 1FN9
1FNN 1FOC 1FP3 1FQT 1FS5 1FSG 1FUU 1G61 1G71 1G8Q 1GEQ 1GG4 1GGG 1GHE 1GIQ
1GL4 1GNW 1GNX 1GOI 1GPE 1GQA 1GTD 1GU2 1GU7 1GUD 1GVE 1GVF 1GVK 1GVU 1GXM
1GYG 1GYO 1H03 1H1O 1H2B 1H32 1H3F 1H3L 1H4P 1H4R 1H4X 1H6G 1H7S 1H80 1H8G
1H8P 1H97 1H9O 1HEK 1HKQ 1HLC 1HPL 1HRU 1HSL 1HST 1HY5 1I19 1I31 1I3Z 1I4J
1I4N 1I4U 1I7K 1IHN 1II2 1IJY 1IN0 1IO7 1IOO 1IPS 1IQ4 1IRD 1IRX 1ISI 1IT2
1ITH 1ITV 1IWM 1IX9 1IXC 1IYB 1IZ5 1J0W 1J1N 1J2X 1J3M 1J6R 1J71 1J7J 1J83
1JAT 1JEK 1JET 1JFL 1JH6 1JHF 1JI1 1JIH 1JL0 1JL9 1JMK 1JMT 1JO0 1JR2 1JS8
1JVN 1JYA 1K07 1K0E 1K38 1K3S 1K4Z 1K66 1K68 1K6D 1K8Q 1KAG 1KAP 1KCF 1KHV
1KJN 1KMT 1KNQ 1KOL 1KPT 1KRH 1KU1 1KUG 1KUT 1KWA 1KXI 1KXJ 1KYF 1L5J 1L6R
1L7A 1L7M 1L8R 1L9M 1LB6 1LEH 1LF6 1LK0 1LKK 1LM4 1LM5 1LM7 1LNZ 1LQT 1LWJ
1LXD 1LYQ 1M0Z 1M1F 1M1Z 1M2D 1M45 1M48 1M55 1M6U 1M8A 1MBY 1MI1 1MIW 1MJH
1MK4 1MKI 1MKZ 1MOL 1MPG 1MQS 1MQV 1MY7 1MZG 1N08 1N0S 1N1B 1N2Z 1N45 1N46
1N7H 1N8V 1NBQ 1NCN 1ND4 1NNW 1NO7 1NOW 1NPE 1NQ7 1NQJ 1NS5 1NSZ 1NTV 1NU0
1NU4 1NUB 1NUL 1NUU 1NXM 1O0W 1O12 1O5U 1O63 1O7I 1O81 1O8B 1OAI 1OBB 1OBO
1OBX 1OCU 1ODZ 1OF3 1OFZ 1OH0 1OHU 1OIZ 1OJ5 1OMZ 1ON2 1OOH 1OQJ 1OW4 1P0K
1P1X 1P4U 1P5T 1P7W 1P9L 1P9Y 1PAM 1PBW 1PD3 1PE9 1PFB 1PGU 1PKH 1PP3 1PP4
1PQ4 1PQH 1PS1 1PT6 1PUI 1PX5 1PXY 1PZL 1PZX 1Q1A 1Q3O 1Q67 1Q77 1Q7F 1Q8Y
1QAH 1QDL 1QEX 1QF8 1QFT 1QGR 1QH5 1QJC 1QJJ 1QJS 1QKR 1QKS 1QLS 1QO2 1QOZ
1QSD 1QUP 1QW9 1QWT 1QYA 1QYR 1R12 1R1D 1R77 1R7A 1R7L 1R9D 1RD5 1REG 1RG8
1RHF 1RHY 1RIF 1RKI 1RKQ 1RP0 1RRL 1RRM 1RW0 1RYL 1RZU 1RZX 1S0P 1S4K 1S4N
1S5P 1S98 1S9R 1SEI 1SFD 1SFL 1SH0 1SH8 1SJ1 1SMO 1SMX 1SQJ 1SQU 1SUL 1SW6
1SWV 1SZ0 1SZH 1SZW 1T0I 1T0P 1T1V 1T2L 1T3G 1T4O 1T6F 1T6T 1T7R 1T92 1TBX
1TDQ 1TE2 1TE5 1TH8 1THT 1TIQ 1TL9 1TLT 1TOA 1TR8 1TVF 1TVN 1TW4 1U00 1U07
1U19 1U5K 1U5U 1U7B 1UAX 1UC7 1UCG 1UCR 1UEB 1UG3 1UJ2 1UJN 1UJW 1UKC 1UMU
1UMZ 1UOC 1UPK 1UPS 1UQT 1URH 1URJ 1URS 1UTI 1UV7 1UWW 1UWZ 1UXZ 1UZ3 1V1A

1V1P 1V37 1V47 1V74 1V8H 1V96 1V9K 1VA6 1VBK 1VC1 1VC4 1VCD 1VDR 1VDW 1VH5
1VHX 1VI2 1VIA 1VIO 1VJ7 1VJL 1VJQ 1VJU 1VKI 1VL4 1VM7 1VMA 1VMO 1VP2 1VPV
1VQQ 1VQU 1VS3 1VYB 1VZY 1W32 1W5R 1W94 1W9C 1W9P 1W9S 1WB4 1WB7 1WC3 1WDU
1WDV 1WEH 1WKO 1WKR 1WLG 1WMH 1WMS 1WMX 1WN1 1WOQ 1WPN 1WQ6 1WR8 1WRA 1WSC
1WSR 1WUF 1WV2 1WVG 1WWL 1WWM 1WWP 1WZ9 1WZD 1X2I 1X6I 1X7O 1X9Z 1XAH 1XCR
1XFS 1XG2 1XGS 1XHK 1XI3 1XIY 1XJU 1XK9 1XM7 1XM8 1XOC 1XOF 1XQA 1XQR 1XRP
1XRS 1XSZ 1XTN 1XVI 1XVS 1XVW 1XYZ 1XZO 1Y0U 1Y1M 1Y1P 1Y3T 1Y44 1Y4T 1Y5H
1Y71 1Y7Y 1Y9Z 1YAC 1YBX 1YC0 1YC5 1YCD 1YDY 1YF2 1YGA 1YLM 1YLQ 1YLX 1YMT
1YNP 1YOC 1YOD 1YOZ 1YPF 1YPQ 1YPY 1YQ1 1YQ5 1YQD 1YQH 1YRK 1YRR 1YZ4 1YZH
1YZY 1Z1Y 1Z2W 1Z2Z 1Z3E 1Z6U 1Z72 1Z85 1Z96 1ZB1 1ZC6 1ZEE 1ZH8 1ZHH 1ZJ8
1ZKC 1ZKD 1ZKI 1ZLP 1ZPL 1ZQ9 1ZSO 1ZTD 1ZUO 1ZUY 1ZVT 1ZY4 1ZY7 1ZYS 1ZZW
2A0S 2A1K 2A2M 2A2R 2A35 2A5L 2A6A 2A6P 2A70 2A8N 2A9D 2AB5 2ABQ 2ABW 2ACV
2AE2 2AEE 2AFB 2AFC 2AFW 2AG4 2AHF 2AHX 2AIB 2AJA 2AMX 2ANX 2APO 2AQ6 2AQP
2AR0 2ARC 2AS9 2ASU 2AUW 2AVN 2AYT 2AZ4 2B0R 2B1L 2B2N 2B3R 2B3Y 2B4M 2B6C
2B82 2B8N 2B97 2B9D 2B9H 2B9R 2BBA 2BCO 2BGH 2BHG 2BJD 2BJN 2BKL 2BKM 2BLF
2BLN 2BM5 2BON 2BPH 2BPO 2BPS 2BRW 2BRY 2BSJ 2BT6 2BU3 2BV4 2BVF 2BWF 2BWR
2BYC 2BZ9 2C0G 2C3I 2C40 2C5U 2C77 2C8J 2C95 2CAR 2CAY 2CB8 2CC0 2CC3 2CFA
2CFO 2CGK 2CI5 2CIA 2CJ4 2CJP 2CKD 2CN3 2CO5 2CU3 2CUN 2CV8 2CVH 2CVI 2CX6
2CX7 2CXD 2CY9 2D1G 2D1H 2D42 2D4G 2D5C 2DB0 2DB7 2DBS 2DC0 2DC3 2DC4 2DEB
2DFJ 2DFY 2DI4 2DOK 2DPR 2DPS 2DPY 2DQ4 2DQA 2DQL 2DQW 2DS5 2DSJ 2DTC 2DUR
2DXU 2DYJ 2E12 2E2E 2E3P 2E5Y 2E85 2E8G 2E8Y 2EAB 2EAV 2EAY 2EBE 2EBJ 2EEN
2EF8 2EG4 2EGG 2EGJ 2EGZ 2EHP 2EIH 2EIS 2EIX 2EJA 2EJN 2EJQ 2EKC 2ERV 2EUC
2EX0 2EXV 2F20 2F23 2F25 2F31 2F37 2F3O 2F4E 2F4M 2F51 2F5J 2F5Y 2F7L 2F8M
2F8Y 2F9H 2F9S 2FAE 2FAO 2FAZ 2FCO 2FCT 2FCW 2FEA 2FFG 2FFI 2FFU 2FH5 2FHP
2FHQ 2FHZ 2FIA 2FJR 2FK5 2FLU 2FN0 2FNA 2FNO 2FP1 2FPR 2FSH 2FSK 2FT0 2FTR
2FTX 2FU4 2FV7 2FVU 2FYX 2FZF 2G09 2G3W 2G58 2G6T 2G7Z 2G8L 2GA1 2GAI 2GAK
2GCL 2GCO 2GD9 2GDQ 2GEC 2GF3 2GF4 2GFF 2GGS 2GGZ 2GHA 2GHV 2GIY 2GJ3 2GLZ
2GMF 2GMQ 2GN4 2GOM 2GOP 2GP4 2GPY 2GPZ 2GRR 2GRU 2GS9 2GSO 2GSV 2GT1 2GV9
2GVY 2GZ6 2GZB 2GZX 2H1C 2H1E 2H1Y 2H34 2H3H 2H7Z 2H98 2HAL 2HB0 2HBA 2HDI
2HDV 2HEK 2HEV 2HF1 2HF2 2HF9 2HFS 2HI0 2HIH 2HIN 2HJ3 2HJV 2HKE 2HLC 2HLS
2HNL 2HP4 2HPL 2HQ4 2HQ9 2HQY 2HRA 2HRV 2HSI 2HTA 2HU9 2HW6 2HWY 2I02 2I0E
2I1S 2I1Y 2I27 2I2O 2I4R 2I4S 2I58 2I5G 2I6H 2I6K 2I6L 2I74 2I9X 2IA1 2IAB
2IB0 2IBN 2IC2 2ICH 2ID1 2IDL 2IEP 2IEW 2IG3 2IM8 2IMZ 2IN5 2INW 2IQJ 2IRP
2IRU 2ISM 2ITB 2ITM 2IUY 2IXN 2IXO 2IXS 2IYG 2IYK 2IZ6 2J16 2J1V 2J4D 2J5B
2J5Y 2J8I 2J9W 2JBV 2JBX 2JCB 2JD4 2JDA 2JDJ 2JE8 2JEM 2JEP 2JF7 2JFZ 2JGB
2JHN 2JIG 2JIK 2JJ7 2JK9 2JKG 2JKH 2MSB 2NLI 2NLV 2NOG 2NRV 2NS9 2NTE 2NTT
2NTX 2NUJ 2NV0 2NW0 2NYU 2NZ5 2O16 2O1E 2O1K 2O1Q 2O2K 2O2T 2O30 2O3B 2O3I
2O5A 2O5H 2O5N 2O62 2O6L 2O6P 2O7G 2O8S 2OA9 2OAF 2OB3 2OB9 2OD0 2OD4 2ODA
2ODM 2OEE 2OER 2OFC 2OFP 2OFY 2OG1 2OGI 2OJL 2OKC 2OKG 2OL7 2OLW 2OM6 2OOC
2OOI 2OPI 2OQ1 2OQA 2OQB 2OQC 2OQQ 2ORV 2ORW 2OTN 2OUS 2OVS 2OWA 2OWL 2OXC
2OXL 2OY9 2OYK 2OZ5 2OZJ 2OZV 2OZZ 2P08 2P0M 2P11 2P12 2P13 2P1A 2P1G 2P35
2P38 2P3P 2P4P 2P4Z 2P62 2P6C 2P6H 2P6X 2P7I 2P8J 2P9R 2PA2 2PBF 2PD2 2PF6
2PFI 2PHK 2PIE 2PIF 2PK3 2PKE 2PKF 2PLG 2PLR 2PMA 2PNZ 2POF 2PPT 2PPW 2PQG
2PQV 2PR7 2PR8 2PRV 2PRX 2PSP 2PW0 2PX6 2PYG 2PZ0 2PZE 2PZI 2Q03 2Q0N 2Q24
2Q2B 2Q2G 2Q3F 2Q3G 2Q3X 2Q5C 2Q5W 2Q6O 2Q7T 2Q7X 2Q83 2Q8X 2Q9O 2QA9 2QAI
2QB7 2QCQ 2QCU 2QCX 2QDQ 2QDR 2QE8 2QEB 2QF4 2QF9 2QG3 2QH5 2QH9 2QHQ 2QJ3
2QJ8 2QJV 2QJZ 2QKH 2QKL 2QMW 2QN4 2QND 2QOS 2QRR 2QS8 2QSJ 2QSQ 2QSX 2QTY
2QV0 2QV5 2QX5 2QXX 2QXY 2QY1 2QY6 2QYC 2QYV 2QZ7 2QZA 2QZC 2R15 2R19 2R25

2R2A 2R5X 2R6J 2R6O 2R6Z 2R76 2R85 2R8B 2R8Q 2R8R 2RA4 2RAD 2RB6 2RBD 2RBG
2REE 2REK 2RFM 2RG4 2RG8 2RI9 2RJI 2RJW 2RKK 2RL8 2RMP 2SCP 2SQC 2UVF 2UWI
2UXT 2V1Q 2V1Y 2V25 2V27 2V2F 2V33 2V3T 2V3Z 2V5C 2V5E 2V6U 2V6V 2V8P 2V94
2V9B 2V9T 2VA8 2VCY 2VD3 2VE3 2VGX 2VH1 2VH3 2VHA 2VHF 2VK7 2VKP 2VLQ 2VLU
2VNG 2VOB 2VOK 2VOZ 2VP8 2VPN 2VPP 2VPV 2VQ3 2VQH 2VQQ 2VRN 2VVE 2VVG 2VVW
2VXB 2VXG 2VYI 2VZC 2W00 2W1J 2W1K 2W2G 2W3G 2W3Y 2W50 2W53 2W56 2W59 2W5F
2W5Z 2W7A 2W7V 2W7Z 2W8D 2W8M 2W9J 2W9T 2W9X 2WB7 2WB9 2WCR 2WD6 2WE8 2WEE
2WEK 2WFH 2WFV 2WG7 2WHN 2WIV 2WJ9 2WKF 2WNH 2WNY 2WOD 2WOK 2WP4 2WPX 2WRZ
2WTP 2WUQ 2WVQ 2WZ1 2X02 2X03 2X0K 2X1Q 2X32 2X3J 2X4D 2X4K 2X5Q 2X61 2X6R
2X7X 2X8S 2X98 2X9J 2X9Q 2XCJ 2XE4 2XEP 2XES 2XET 2XEX 2XFA 2XFV 2XGG 2XGU
2XHA 2XHF 2XHS 2XI8 2XI9 2XMJ 2XMO 2XMX 2XOC 2XOL 2XOT 2XPP 2XQX 2XR1 2XSS
2XSW 2XT2 2XTL 2XTM 2XTY 2XUA 2XUS 2XVC 2XVM 2XXN 2XYI 2XZ4 2XZ8 2XZI 2Y1H
2Y2X 2Y43 2Y4J 2Y7E 2Y7I 2Y7S 2Y8E 2Y8U 2Y9M 2YB7 2YCH 2YEQ 2YFQ 2YG2 2YHN
2YJ6 2YJG 2YKT 2YMY 2YN1 2YN5 2YN7 2YNA 2YOA 2YOC 2YOR 2YQY 2YQZ 2YR1 2YV9
2YVR 2YWW 2YXD 2YXE 2YXO 2YXW 2YY6 2YYB 2YYS 2YYV 2Z0U 2Z22 2Z26 2Z5B 2Z5D
2Z64 2Z73 2Z8F 2Z8G 2ZAY 2ZBI 2ZC2 2ZCA 2ZFU 2ZGY 2ZKT 2ZMV 2ZOS 2ZOU 2ZSI
2ZTB 2ZU9 2ZVD 2ZVR 2ZWA 2ZWI 2ZWR 2ZX2 2ZXD 2ZYR 2ZZ8 2ZZV 3A07 3A0Y 3A1D
3A1S 3A21 3A24 3A35 3A43 3A45 3A4M 3A4R 3A4T 3A54 3A5I 3A6S 3A9F 3A9L 3AAG
3AAY 3AB1 3ABG 3ADR 3AEH 3AEI 3AFF 3AFM 3AGX 3AHN 3AIH 3AJ6 3AJA 3AJR 3AKJ
3AL3 3AMI 3AMN 3ANW 3AOF 3APQ 3APR 3APT 3APU 3APZ 3AQ9 3AQG 3AQL 3AS5 3ASL
3ATY 3AU4 3AVR 3AWU 3AXA 3AXD 3AYC 3AZD 3AZO 3B0F 3B0P 3B4N 3B4Q 3B5E 3B5I
3B5Q 3B6H 3B73 3B7S 3B85 3BA3 3BBD 3BBZ 3BDV 3BE3 3BEU 3BF7 3BFV 3BGA 3BGE
3BGH 3BGY 3BH4 3BHD 3BHW 3BIT 3BJ4 3BMX 3BNW 3BO6 3BOH 3BOO 3BP3 3BQ9 3BQO
3BQP 3BRN 3BRS 3BS6 3BS7 3BTP 3BUS 3BVO 3BVP 3BW1 3BWS 3BWV 3BXP 3BXW 3BYP
3BZB 3BZY 3C0G 3C0U 3C1A 3C3R 3C4N 3C4S 3C4V 3C57 3C5N 3C7M 3C8C 3C8L 3C9F
3C9G 3C9H 3C9Q 3CB2 3CBW 3CEG 3CEI 3CEU 3CEX 3CFU 3CG6 3CG7 3CHH 3CIO 3CIT
3CJP 3CK1 3CKC 3CLK 3CNH 3CNR 3CNY 3COB 3COK 3COL 3COV 3CP7 3CPT 3CQB 3CQC
3CQL 3CRN 3CT6 3CU2 3CU5 3CUC 3CV0 3CWC 3CWF 3CWV 3CX3 3CYG 3CZ1 3CZB 3CZH
3D21 3D34 3D37 3D3B 3D3Q 3D4J 3D59 3D5J 3D5P 3D6I 3D6R 3D6W 3D7A 3D8C 3D8D
3D8U 3D9Y 3DA5 3DAD 3DB0 3DBA 3DBG 3DC6 3DCD 3DDE 3DDL 3DEP 3DEU 3DGP 3DKA
3DLQ 3DME 3DNF 3DNS 3DNT 3DO8 3DOH 3DR2 3DRF 3DRN 3DRW 3DS2 3DSK 3DTB 3DTN
3DUP 3DWM 3DWV 3DXO 3DXQ 3DXR 3DYJ 3DYN 3DZV 3E0X 3E11 3E2V 3E3R 3E48 3E4W
3E57 3E58 3E7H 3E7J 3E9C 3E9G 3EA0 3EAE 3EAG 3EB8 3EB9 3EC3 3EC4 3EC9 3ECN
3ECQ 3ECR 3EDN 3EDP 3EDV 3EE6 3EEA 3EEF 3EEQ 3EFP 3EGR 3EHD 3EIP 3EJW 3ELK
3EMX 3EN9 3ENC 3ENP 3EO6 3EOP 3EOQ 3EOZ 3EP0 3EPS 3EQX 3EQZ 3ERP 3ERX 3ESA
3ESL 3ETC 3ETO 3ETQ 3ETZ 3EU7 3EUS 3EVI 3EVY 3EWI 3EWL 3EWM 3EWO 3EYY 3EZH
3F08 3F0P 3F13 3F1P 3F42 3F4A 3F5H 3F66 3F69 3F6C 3F6I 3F6K 3F6O 3F70 3F7E
3F7Q 3F8B 3F95 3F9U 3FB9 3FBG 3FCD 3FCG 3FCM 3FD4 3FD7 3FDI 3FDW 3FDX 3FE3
3FE4 3FF1 3FF5 3FG7 3FGV 3FHW 3FID 3FIL 3FJV 3FLA 3FLE 3FLT 3FM2 3FM3 3FN1
3FN5 3FNC 3FO3 3FO5 3FPK 3FPN 3FPR 3FQD 3FQM 3FSO 3FUT 3FVD 3FVV 3FVW 3FW3
3FYF 3FZY 3G12 3G1E 3G1J 3G1P 3G23 3G2F 3G2M 3G3R 3G3S 3G46 3G48 3G4D 3G4E
3G5J 3G68 3G8K 3GAE 3GAX 3GAZ 3GBV 3GBY 3GD4 3GDI 3GF5 3GF6 3GFF 3GFV 3GGN
3GHD 3GI7 3GID 3GJ0 3GKN 3GKX 3GLV 3GME 3GMG 3GNL 3GO6 3GOC 3GPK 3GPV 3GQS
3GRA 3GRD 3GRI 3GRN 3GRO 3GRZ 3GUD 3GUE 3GUU 3GV4 3GVE 3GWB 3GWL 3GWR 3GXH
3GYC 3GYZ 3GZ5 3GZA 3H05 3H1Q 3H2B 3H2S 3H30 3H3A 3H3N 3H5J 3H5L 3H7J 3H7O
3H8Q 3H8V 3HA2 3HAM 3HBW 3HCS 3HCW 3HCY 3HDF 3HDT 3HEB 3HFH 3HGT 3HHF 3HHI
3HIS 3HJ4 3HJ6 3HJG 3HK0 3HKL 3HKS 3HKV 3HL1 3HL6 3HLK 3HM4 3HME 3HMT 3HN0
3HN5 3HO6 3HOB 3HPE 3HPK 3HQR 3HRQ 3HRS 3HS3 3HU5 3HV1 3HV2 3HWJ 3HWO 3HWP

3HY0 3HYJ 3I00 3I1A 3I1I 3I3Q 3I3W 3I41 3I4O 3I57 3I5O 3I5R 3I5W 3I6D 3I6S
3I7J 3I83 3I8N 3I9F 3IA1 3IA8 3IAU 3IB3 3IBW 3IBX 3IC5 3ICY 3ID9 3IDF 3IE4
3IE5 3IEG 3IGE 3IHS 3IHT 3IHV 3IIC 3IJL 3IJM 3IJW 3IKB 3INO 3IO1 3IOL 3IPF
3IPJ 3IPO 3IQ0 3IQ2 3IQC 3IQU 3IR9 3IRB 3IS6 3ITE 3ITQ 3ITW 3IU1 3IUK 3IUO
3IUP 3IUS 3IUW 3IUY 3IV7 3IVL 3IVV 3IWF 3IWG 3IX1 3IX3 3IX7 3IX9 3JQ1 3JR7
3JRR 3JRU 3JSB 3JSL 3JT0 3JTN 3JUU 3JW8 3JWI 3JXF 3JXO 3K0Z 3K1R 3K1W 3K2A
3K2N 3K2O 3K2Z 3K51 3K5O 3K6F 3K6O 3K7B 3K85 3K8G 3K8R 3K9V 3KA5 3KB1 3KB2
3KBQ 3KBY 3KD3 3KD4 3KD6 3KDG 3KEA 3KEP 3KEW 3KEZ 3KF6 3KFA 3KG8 3KG9 3KGK
3KHE 3KHN 3KK7 3KKS 3KLQ 3KM5 3KMA 3KMI 3KMR 3KNB 3KPE 3KS9 3KSM 3KSU 3KTZ
3KUZ 3KWR 3KY9 3KYJ 3KZY 3L01 3L0Q 3L0R 3L12 3L15 3L18 3L32 3L46 3L50 3L6I
3L6T 3L6V 3L6X 3L7O 3L81 3L8C 3L8E 3L8M 3L9J 3LAE 3LAG 3LAZ 3LB2 3LET 3LF5
3LFR 3LFT 3LG3 3LGB 3LHN 3LHX 3LID 3LIF 3LIU 3LJB 3LJS 3LKB 3LKL 3LLM 3LLP
3LLZ 3LM2 3LMH 3LMN 3LNN 3LNY 3LQ9 3LQM 3LS1 3LS8 3LST 3LUO 3LVC 3LWE 3LX4
3LX6 3LXQ 3LY0 3LYP 3LYX 3M33 3M6Z 3M7A 3M8J 3M8T 3MAB 3MAL 3MAZ 3MB4 3MBC
3MC9 3MCA 3MCB 3MCF 3MCS 3MCW 3MD1 3MD9 3MDF 3ME4 3ME7 3MEA 3MER 3MFD 3MG1
3MGD 3MGG 3MH9 3MIT 3MIZ 3MJQ 3MK4 3MKL 3MNL 3MOZ 3MPC 3MPD 3MQ2 3MR0 3MTI
3MTK 3MTR 3MUQ 3MUX 3MVC 3MVP 3MWB 3MWX 3MX3 3MXO 3MYU 3MYV 3MYX 3MZ2 3N01
3N08 3N10 3N1E 3N4I 3N6Y 3N72 3N89 3N9B 3N9V 3NA5 3NBC 3NCE 3NCX 3NDA 3NDO
3NEK 3NEQ 3NFH 3NFQ 3NGF 3NHM 3NI7 3NIQ 3NJ2 3NJE 3NK6 3NKL 3NKU 3NME 3NMW
3NNG 3NNN 3NNS 3NO8 3NPF 3NPP 3NQN 3NR1 3NRF 3NRH 3NRL 3NRX 3NT8 3NTK 3NTX
3NUF 3NW0 3NWP 3NY3 3NYI 3NZE 3NZN 3NZZ 3O0A 3O0L 3O0Q 3O0X 3O14 3O2U 3O53
3O5Y 3O60 3O66 3O6W 3O7A 3O83 3O8Q 3OAJ 3OBE 3OBF 3OBH 3OBL 3OBQ 3OBY 3OCO
3OCP 3OG5 3OG6 3OG7 3OGN 3OHE 3OIQ 3OKW 3OKZ 3OL3 3OMD 3OMT 3ON3 3ON9 3ONM
3OOV 3OOX 3OP6 3OPE 3OQI 3OT2 3OTN 3OVP 3OWC 3OWG 3OXP 3OY2 3OYO 3OYY 3OZD
3OZI 3OZX 3P09 3P0U 3P1U 3P2E 3P3Q 3P3V 3P5R 3P69 3P6A 3P9X 3PAF 3PC6 3PD7
3PE5 3PES 3PET 3PF8 3PG7 3PGG 3PGS 3PH9 3PHG 3PHX 3PIV 3PJP 3PM6 3PMC 3PMG
3PNR 3PQH 3PQU 3PRB 3PSM 3PSQ 3PT3 3PT8 3PU8 3PVE 3PWX 3Q18 3Q1I 3Q2J 3Q49
3Q6K 3Q6V 3Q72 3Q7R 3Q87 3QAT 3QAX 3QB8 3QC2 3QC4 3QE2 3QEE 3QEK 3QF2 3QHB
3QHP 3QHQ 3QI7 3QIJ 3QIS 3QN9 3QPI 3QR2 3QR7 3QRC 3QSL 3QSZ 3QT5 3QTA 3QTG
3QTM 3QU5 3QUF 3QVL 3QVM 3QW9 3QWG 3QX1 3QYE 3QYY 3QZ4 3QZM 3QZR 3R07 3R0J
3R15 3R1J 3R27 3R41 3R42 3R4R 3R4S 3R5Z 3R62 3R6A 3R7A 3R7G 3RA5 3RAO 3RAU
3RB5 3RBY 3RC4 3RDK 3RE1 3RE4 3RFS 3RGC 3RGH 3RH0 3RHY 3RHZ 3RI0 3RJT 3RK1
3RKC 3RLS 3RMH 3RNQ 3RO3 3ROI 3ROT 3RP2 3RPJ 3RQ9 3RS1 3RUX 3RV6 3RY0 3RY3
3S0R 3S0T 3S2X 3S4K 3S5B 3S5F 3S5W 3S63 3S6E 3S7D 3S8I 3S8K 3S8P 3S93 3S95
3S9U 3SAF 3SAO 3SCZ 3SD4 3SEI 3SEO 3SFW 3SG8 3SGH 3SHP 3SIM 3SIT 3SJ5 3SKV
3SLU 3SLZ 3SO6 3SOJ 3SOK 3SON 3SOV 3SP1 3SP4 3SPE 3SQF 3SQJ 3SRI 3STY 3SUB
3SUK 3SWH 3SYL 3SZ6 3T0P 3T13 3T1O 3T47 3T4L 3T5G 3T5X 3T6K 3T8B 3T9G 3T9K
3TB6 3TBH 3TC8 3TCA 3TCN 3TCR 3TCV 3TDN 3TDQ 3TDV 3TE8 3TEB 3TEJ 3TEK 3TEV
3TFG 3TII 3TIQ 3TKF 3TL1 3TLQ 3TM8 3TOD 3TOV 3TP2 3TP9 3TQF 3TQW 3TRB 3TSA
3TSJ 3TSM 3TTM 3TUF 3TV1 3TVA 3TVT 3TWD 3TWE 3TWF 3TWK 3TX3 3TYQ 3TZG 3TZT
3U0H 3U0J 3U1D 3U1U 3U1X 3U21 3U23 3U3B 3U4T 3U4Y 3U4Z 3U7R 3U7Z 3U80 3U8V
3U96 3U9J 3U9Q 3UAN 3UC4 3UEC 3UES 3UGF 3UHA 3UID 3UIW 3UL3 3ULJ 3ULL 3ULT
3ULY 3UMZ 3UN7 3UO3 3UP1 3UP3 3UPV 3UR8 3URR 3USH 3USS 3USY 3UT4 3UUG 3UV0
3UV1 3UXN 3UY7 3UYJ 3V0D 3V1E 3V30 3V33 3V3L 3V43 3V48 3V67 3V69 3V8D 3V8I
3V97 3V98 3VAS 3VAY 3VCC 3VCF 3VDH 3VEJ 3VF1 3VFZ 3VHS 3VJA 3VJE 3VJP 3VK5
3VKG 3VMT 3VO2 3VOQ 3VPP 3VPS 3VRC 3VTA 3VTH 3VTX 3VU2 3VU4 3VU9 3VUP 3VUS
3VV1 3VV3 3VV5 3VX3 3VX4 3VYP 3VZI 3W08 3W0E 3W0K 3W19 3W1O 3W2Y 3W3W 3W4S
3W57 3W5F 3W5S 3W6P 3W7T 3W9S 3W9V 3WA4 3WA8 3WAE 3WAS 3WDF 3WDW 3WE2 3WE5

3WEA 3WEU 3WFI 3WH9 3WHT 3WI7 3WJ9 3WKY 3WL2 3WL4 3WL6 3WMD 3WMG 3WMI 3WNO
3WOL 3WPW 3WQO 3WUR 3WV4 3WWN 3WX1 3WYD 3ZBD 3ZBO 3ZD2 3ZFI 3ZG6 3ZGJ 3ZH5
3ZHO 3ZIH 3ZIL 3ZIT 3ZIU 3ZJE 3ZK9 3ZL1 3ZME 3ZMR 3ZO9 3ZPY 3ZQS 3ZRG 3ZTP
3ZWF 3ZXC 3ZXF 3ZXN 3ZY7 3ZYG 3ZYL 3ZYR 3ZYW 4A0E 4A0Z 4A2O 4A37 4A48 4A6F
4A6V 4A7U 4A7W 4A8H 4AAZ 4ACV 4ADN 4ADT 4ADY 4ADZ 4AE4 4AEE 4AEF 4AGG 4AHC
4AJW 4AKL 4AKM 4ALF 4AM6 4AMJ 4APX 4AQN 4ARV 4ASR 4AU9 4AUC 4AUP 4AVB 4AVR
4AWX 4AXK 4AXN 4AY0 4AYA 4AYG 4B0Z 4B1Y 4B2N 4B3B 4B45 4B5Q 4B61 4B6G 4B6M
4B6X 4B8B 4B8E 4B91 4B93 4B9G 4BD2 4BEG 4BFA 4BG2 4BGO 4BHR 4BI3 4BLU 4BND
4BOP 4BPG 4BPZ 4BQ4 4BQ9 4BQN 4BQU 4BRC 4BS6 4BSZ 4BUC 4BUU 4BVQ 4BVX 4BWO
4BWV 4BX8 4BXH 4C0R 4C16 4C1D 4C1L 4C1S 4C23 4C29 4C3D 4C76 4C7A 4C7D 4C8B
4C97 4C9Y 4CA1 4CB7 4CBP 4CDJ 4CEM 4CGR 4CGS 4CGY 4CHF 4CHH 4CI7 4CI8 4CJ0
4CJ9 4CK4 4CMR 4CQ8 4CRW 4CSD 4CU9 4CUA 4CXF 4CXV 4CZJ 4CZX 4D05 4D0O 4D0Y
4D2C 4D2O 4D8I 4D9I 4D9S 4DBG 4DCB 4DCZ 4DEY 4DGF 4DGH 4DHK 4DI8 4DIX 4DJB
4DJG 4DKC 4DKN 4DLH 4DLQ 4DM4 4DO4 4DO7 4DOI 4DOK 4DOO 4DOV 4DQ9 4DQZ 4DS2
4DSD 4DT5 4DTE 4DY0 4DYH 4DYN 4DYW 4DZM 4DZZ 4E15 4E19 4E1Y 4E3Y 4E57 4E5V
4E5W 4E6F 4E7S 4E8U 4E94 4E9J 4EBR 4ECO 4EDH 4EE6 4EEI 4EET 4EF0 4EFO 4EG0
4EGD 4EH1 4EHS 4EHU 4EI0 4EI7 4EIB 4EIR 4EIS 4EIV 4EJR 4EMT 4EP4 4EPP 4EQB
4EQQ 4ERC 4ERY 4ES8 4ETV 4ETZ 4EUK 4EUU 4EVQ 4EVU 4EVW 4EW5 4EWI 4EWL 4EYG
4EYZ 4EZG 4F0D 4F14 4F1J 4F27 4F3V 4F3Y 4F44 4F4F 4F7K 4F7O 4F82 4FCH 4FCZ
4FD4 4FD9 4FDI 4FDX 4FDY 4FEK 4FET 4FGQ 4FHR 4FID 4FKB 4FKZ 4FP1 4FPW 4FQ5
4FQD 4FRF 4FRX 4FXQ 4FYP 4FYT 4FZL 4FZP 4FZV 4G0I 4G0M 4G0S 4G1I 4G2B 4G2C
4G2U 4G37 4G3B 4G3C 4G3V 4G4K 4G4L 4G4M 4G6Q 4G6U 4G7X 4G8K 4G9M 4G9S 4GBF
4GBO 4GBS 4GC1 4GCN 4GCS 4GE6 4GEK 4GGG 4GHB 4GIW 4GKC 4GKF 4GKG 4GKM 4GKP
4GL6 4GMN 4GNE 4GNI 4GNS 4GNU 4GOF 4GQ6 4GUC 4GVB 4GVF 4GVO 4GXB 4GXL 4GYT
4H05 4H0A 4H0C 4H0K 4H2D 4H4D 4H5I 4H5S 4H61 4H6Q 4H7X 4H87 4H8F 4H8M 4HAP
4HBQ 4HC8 4HCE 4HCI 4HDH 4HEH 4HEO 4HEQ 4HFS 4HG2 4HH6 4HHV 4HI7 4HI8 4HIA
4HIL 4HJD 4HJZ 4HKE 4HKG 4HL0 4HL2 4HLS 4HN9 4HNE 4HNH 4HP8 4HQZ 4HR1 4HRZ
4HS5 4HSS 4HT3 4HU5 4HW8 4HWU 4HWV 4HY4 4HYJ 4HYL 4HYN 4HZR 4I1K 4I1U 4I2Z
4I3G 4I4K 4I4O 4I5T 4I6P 4I6R 4I82 4I84 4I86 4I93 4IB2 4ID2 4ID3 4IGA 4IGW
4IHE 4IHZ 4IJ5 4IJR 4IJZ 4IKN 4ILO 4ILV 4IMQ 4IN0 4IN9 4INA 4INE 4INO 4INZ
4IO2 4IU3 4IUP 4IX3 4IXA 4IXJ 4IXN 4IYB 4IZB 4IZK 4J05 4J0X 4J1Y 4J2G 4J2K
4J3H 4J5R 4J6O 4J73 4J7Q 4J8B 4J8C 4J8E 4J8S 4J9C 4JBS 4JCH 4JCW 4JDE 4JE1
4JE6 4JEM 4JES 4JF3 4JGG 4JGI 4JGP 4JGW 4JGX 4JIX 4JJ0 4JJH 4JK8 4JLI 4JMD
4JN3 4JOQ 4JPQ 4JQT 4JR6 4JT4 4JUI 4JVU 4JX0 4JXB 4JXD 4JXE 4JY3 4JZP 4JZQ
4JZZ 4K00 4K02 4K05 4K0D 4K12 4K1C 4K28 4K2W 4K35 4K3L 4K4K 4K5A 4K6J 4K7J
4K7K 4K8Y 4K9Q 4KBM 4KCE 4KDX 4KED 4KF8 4KFS 4KFW 4KGD 4KGH 4KH6 4KH7 4KH9
4KHO 4KJM 4KJR 4KMD 4KN8 4KNC 4KNK 4KP2 4KPO 4KQR 4KRG 4KRT 4KT1 4KT3 4KTW
4KUJ 4KUN 4KV2 4KV9 4KWY 4KX8 4KYU 4KYX 4L00 4L0R 4L3N 4L3R 4L3T 4L4W 4L51
4L5G 4L68 4L6S 4L6U 4L7A 4L7X 4L8I 4L9O 4L9U 4LA2 4LAS 4LBA 4LCI 4LE7 4LEB
4LEC 4LIR 4LJI 4LJL 4LK2 4LLD 4LN2 4LN9 4LNL 4LOW 4LP4 4LPS 4LQ8 4LQC 4LQX
4LS4 4LUB 4LV5 4LW8 4LWK 4LXO 4LXQ 4M0H 4M1A 4M1B 4M1Q 4M3O 4M4D 4M8R 4M91
4MAC 4MAE 4MAK 4MAL 4MDU 4ME9 4MES 4MF9 4MG3 4MH1 4MHV 4MIK 4MIX 4MJ2 4MJD
4MJG 4MJK 4MLM 4MLZ 4MM2 4MMG 4MN5 4MN7 4MNW 4MO1 4MOV 4MPB 4MPM 4MPS 4MQB
4MR0 4MTL 4MUV 4MVE 4MW0 4MY6 4MYA 4MYP 4MYV 4MZ3 4MZJ 4MZZ 4N01 4N04 4N06
4N0K 4N0R 4N0V 4N3P 4N3V 4N4U 4N6A 4N6C 4N6F 4N7F 4N7W 4N82 4N8O 4N8Y 4N9Z
4NC7 4NCR 4NE2 4NET 4NFC 4NHB 4NIR 4NJH 4NKT 4NN2 4NOF 4NOH 4NPL 4NQ8 4NSD
4NSV 4NTG 4NTQ 4NWO 4NX8 4NZV 4O1J 4O1S 4O2H 4O2I 4O2T 4O3V 4O42 4O5P 4O71
4O7H 4O7J 4O8V 4O9D 4O9K 4O9S 4OEV 4OF6 4OFK 4OFQ 4OH7 4OHJ 4OK9 4OKE 4OLK

4OLT 4OM7 4OMV 4ON1 4ONW 4ONY 4OO0 4OO4 4OPM 4OTE 4OUC 4OVS 4OVT 4OWI 4OX6
4OZE 4P0J 4P0T 4P2I 4P2L 4P32 4P3F 4P5E 4P5F 4P5N 4P7B 4P7C 4P7O 4P93 4PAB
4PAS 4PE0 4PFZ 4PH8 4PI3 4PIC 4PID 4PIV 4PKC 4PM4 4PMK 4PMO 4PN6 4PO6 4POW
4PQ1 4PQ9 4PR3 4PSF 4PSR 4PTB 4PUI 4PVC 4PXW 4PXY 4PYS 4PZ7 4Q14 4Q2T 4Q3H
4Q4K 4Q53 4Q5G 4Q60 4Q69 4Q6J 4Q6U 4Q7E 4Q7O 4Q7Q 4Q82 4Q88 4Q8L 4Q9A 4Q9B
4Q9T 4Q9W 4QAK 4QAM 4QAN 4QAS 4QBN 4QC6 4QE0 4QF3 4QGO 4QHJ 4QI0 4QJB 4QJI
4QM9 4QMI 4QO2 4QPM 4QPV 4QSE 4QT9 4QUV 4QWO 4QYB 4R01 4R1K 4R1S 4R23 4R7X
4R80 4R86 4R8O 4R8R 4R9X 4RD8 4RHA 4RHP 4RK6 4RK9 4RPC 4RS2 4TKR 4TL1 4TMX
4TQL 4TR6 4TR7 4TY0 4U13 4U4I 4U99 4U9C 4UNU 4UON 4UOP 4UP0 4UQW 4UQY 4URG
4USQ 4UUU

**Supplementary S2.** List of targets (PDB codes) from Dockground Docking Decoy Set 2 used to benchmark native complex ranking (Kundrotas et al., 2018).

1AY7 1BDJ 1BUH 1BUI 1BVN 1CC0 1CLV 1DFJ 1E6J 1EER 1EWY 1F6M 1F80 1F93 1FCC
1FLE 1FLT 1FQ1 1FRT 1G6V 1G73 1GCQ 1GG2 1GLB 1GPQ 1GPW 1HE1 1HJA 1HYR 1I2M
1I4E 1ICF 1IM9 1IS7 1JIW 1JK9 1JTD 1JZD 1K5G 1KGY 1L9J 1LB2 1LFD 1LTX 1LX5
1M63 1MG2 1ML0 1MQ8 1NBF 1NQL 1OC0 1OFU 1PVH 1QAV 1R5I 1R8S 1RPQ 1S1Q 1SJH
1TCO 1TE1 1TMQ 1U0N 1UAD 1UUG 1V5I 1V7P 1VFB 1VG0 1VRS 1W1I 1WEJ 1WQ1 1WRD
1X86 1XK4 1XT9 1Y8R 1Y8X 1YCS 1YU6 1YVB 1Z5Y 1ZLH 2A1T 2A41 2A5D 2A9K 2ABZ
2AQ3 2B4S 2BCG 2BCN 2BOV 2BQ1 2BWE 2C0L 2C2V 2CH4 2D5R 2EJF 2G45 2GJ7 2GRX
2GWF 2HJ9 2HQS 2HRK 2I25 2IDO 2IJO 2IWT 2J12 2NQD 2NXN 2O25 2O2V 2O3B 2O8V
2OOB 2PU9 2QYI 2RII 2SGE 2UY7 2V55 2VDB 2VRR 2WBW 2WY7 2WY8 2X0B 2X9A 2XGY
2XZ1 2YVJ 2ZAE 2ZU0 2ZVN 3A1P 3A8I 3AV0 3BH6 3BK3 3BP8 3BS5 3BX1 3BX7 3CBK
3CII 3CU1 3CVH 3CW2 3D3C 3D5O 3D5R 3DAW 3E2L 3F1P 3F7P 3FAP 3FN1 3FPU 3G3A
3G6D 3GE3 3HI6 3HZI 3K1I 3KLD 3L1Z 3L4Q 3L5N 3L89 3LB8 3LTF 3LVJ 3LWN 3M18
3MJ7 3MZW 3OED 3OJ4 3ONA 3ONL 3OUN 3PNL 3PRO 3PRP 3PV6 3QC8 3QLU 3R66 3R9A
3RJ3 3RNK 3RRM 3S36 3S3X 3S5L 3T1Q 3T5G 3TG1 3UAI 3ULQ 3VLB 3VXU 3W31 3WKT
3WWK 3WWN 3ZN6 3ZNZ 4BI8 4BMP 4BOS 4BOZ 4BWS 4C0O 4C6T 4CPA 4CT4 4CU4 4DI3
4DN4 4DSS 4E2I 4EJX 4EMJ 4ETQ 4ETW 4EXT 4F7G 4FA8 4FI3 4FME 4GED 4GMJ 4GOJ
4HCN 4HH3 4HMY 4HQP 4HRE 4HX3 4I5L 4ICG 4IHF 4ILH 4J38 4J4L 4JAV 4JCV 4JHP
4JO9 4JQW 4JX1 4K0V 4K71 4KEH 4KI1 4KR0 4KXZ 4KYI 4L41 4LHU 4LNU 4LRZ 4LSX
4LW4 4LX0 4M3K 4M7L 4MDK 4MNE 4MRT 4N6O 4NBE 4NIF 4NYI 4NZL 4O4B 4P2A 4P4H
4P4Q 4P69 4PDC 4PJ2 4PJ8 4POU 4PW9 4QD2 4QTI 4R62 4R9Y 4RF0 4RIX 4RWS 4S10
4TXV 4V2C 4WLR 4WM0 4WZA 4XHU 4XIG 4XR8 4XS0 4YEB 4ZFR 5AUP 5BMU 5BVP 5C3I
5C7X 5CEC 5CRA 5CVO 5E0K 5E8D 5EG3

**Supplementary S3.** List of protein complexes (PDB codes) from the 'Bahadur' and 'Duarte' datasets (Bahadur et al., 2004; Duarte et al., 2012) used to benchmark identification of crystal contacts.

**Bahadur dataset:**
13PK 1A7V 1AD5 1AFK 1AG9 1AMU 1ATL 1AW7 1BC2 1BIN 1BKZ 1BYO 1C02 1CKI 1CQX 1DSU 1DYS 1EHY 1EPA 1FGK 1FJM 1FMT 1G2A 1GAR 1ILR 1KPT 1KWA 1MPG 1MSS 1NAW 1QCI 1QDM 1QHA 1QPA 1QTQ 1RB3 1SHK 1THE 1THT 1TOA 1URP 1VBT 1XGS 256B 2ATJ 2BC2 2BLS 2G3P 2J5X 2SCP 2SHP 2TPS 2UGI 3MHT 3PMG 5TSS 830C

**Duarte dataset:**
1CQX 1EJD 1FPO 1J96 1LQT 1N45 1PP3 1TOA 1UEB 1VQQ 1W9Q 1WOQ 1YNQ 1ZLQ 2CKI 2E1V 2F37 2GAS 2IPI 2J46 2OW9 2W20 2WBM 2X26 2YZ1 2ZYR 3C1D 3F0O 3H30 3IRB 3ITA 3LVD 3MG1 3MHJ 3N5C

**Supplementary S4.** Distribution of contact overlap fractions in native complexes used for construction of residue preference matrices. Vertical dashed lines indicate 80th (blue) and 90th (black) percentile of contacts. Our sigmoidal clash penalty function has been optimized such that 80% of the contacts are not penalized (left of blue dashed line), 10% of the contacts are penalized logistically (between blue and black dashed line) and 10% of the worst contacts (right of black dashed line) are assigned a score of 0.

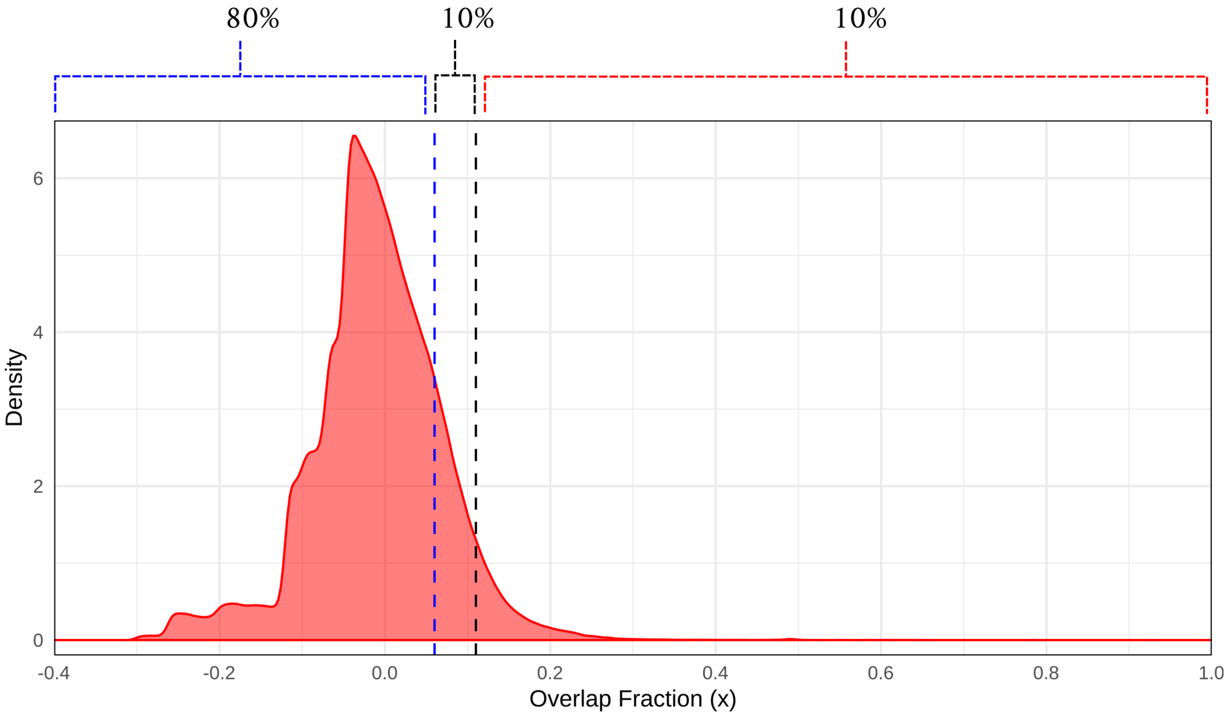

| **Binding mode** | **Target** | **Template** | **Chains** | **Sequence Identity** | **MolPDF** | **DOPE** | **Normalized DOPE** | **GA341** |
|---|---|---|---|---|---|---|---|---|
| AMB7 | AMD9 | 1KXT | A,B | 93.5 | 4190.2 | -71265.7 | -1.25 | 1.00 |
| AMB7 | AMD10 | 1KXT | A,B | 92.7 | 4282.5 | -70850.3 | -1.23 | 1.00 |
| AMD9 | AMB7 | 1KXQ | A,H | 93.8 | 4127.8 | -72921.5 | -1.34 | 1.00 |
| AMD9 | AMD10 | 1KXQ | A,H | 92.3 | 4002.9 | -71715.5 | -1.33 | 1.00 |
| AMD10 | AMB7 | 1KXV | A,C | 93.5 | 4190.2 | -71265.7 | -1.25 | 1.00 |
| AMD10 | AMD9 | 1KXV | A,C | 92.0 | 4058.8 | -71137.2 | -1.23 | 1.00 |

**Supplementary S5.** Statistics of modeling the non-native VHH-PPA binding modes.

**Supplementary S6 (a).** Interacting residue pair atomic propensities at a distance threshold of 4 Å and main chain - main chain mode of interaction.

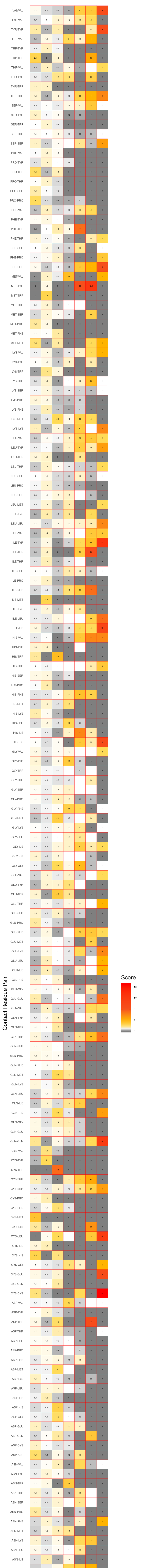

**Supplementary S6 (b).** Interacting residue pair atomic propensities at a distance threshold of 4 Å and side chain - side chain mode of interaction.

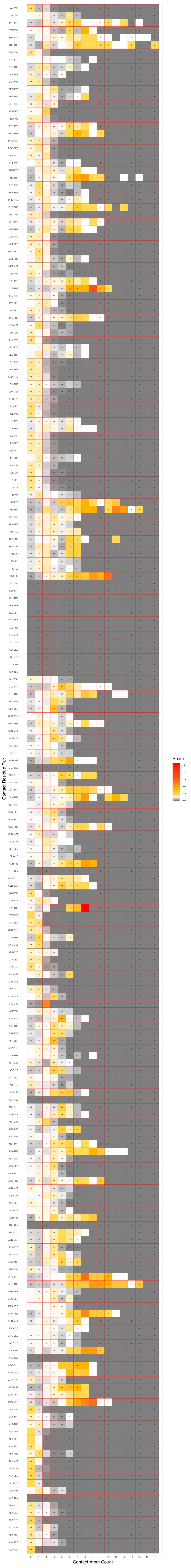

**Supplementary S6 (c).** Interacting residue pair atomic propensities at a distance threshold of 4 Å and main chain - side chain mode of interaction.

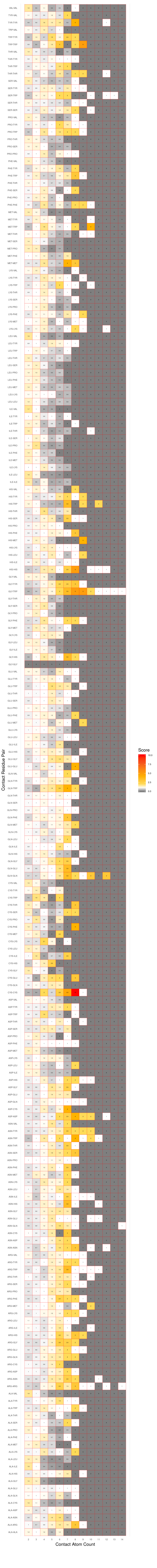

**Supplementary S6 (d).** Interacting residue pair atomic propensities at a distance threshold of 6 Å and main chain - main chain mode of interaction.

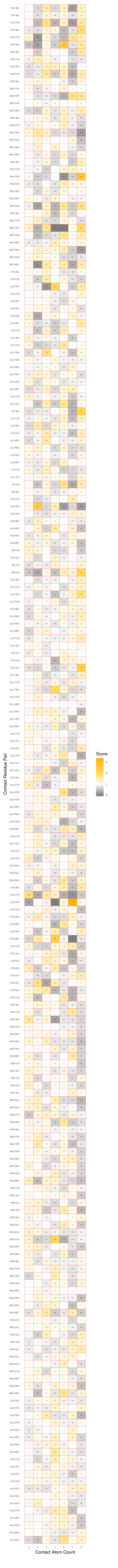

**Supplementary S6 (e).** Interacting residue pair atomic propensities at a distance threshold of 6 Å and side chain - side chain mode of interaction.

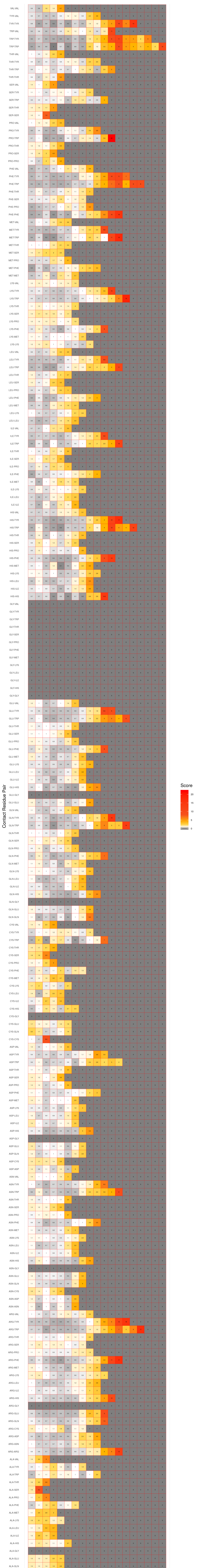

**Supplementary S6 (f).** Interacting residue pair atomic propensities at a distance threshold of 6 Å and main chain - side chain mode of interaction.

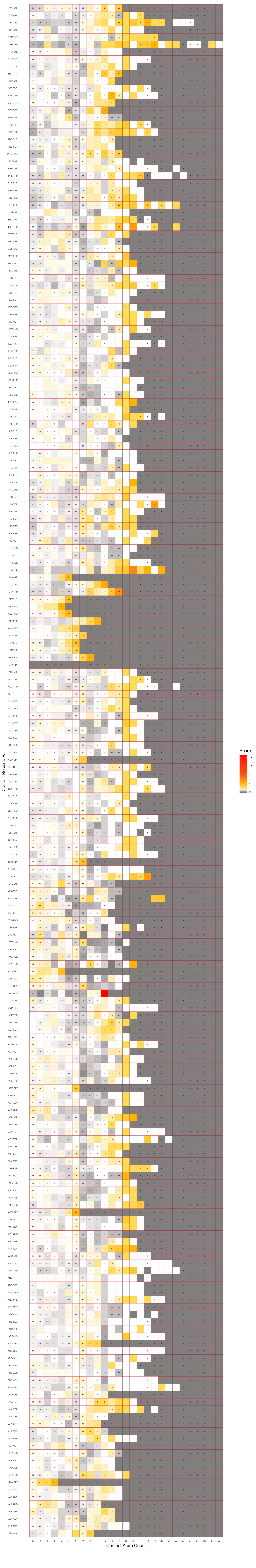

**Supplementary S6 (g).** Interacting residue pair atomic propensities at a distance threshold of 8 Å and main chain - main chain mode of interaction.

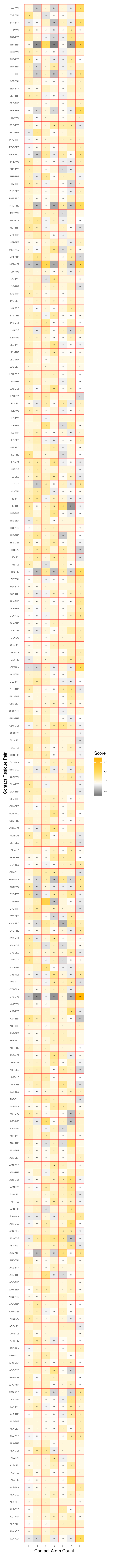

**Supplementary S6 (h).** Interacting residue pair atomic propensities at a distance threshold of 8 Å and side chain - side chain mode of interaction.

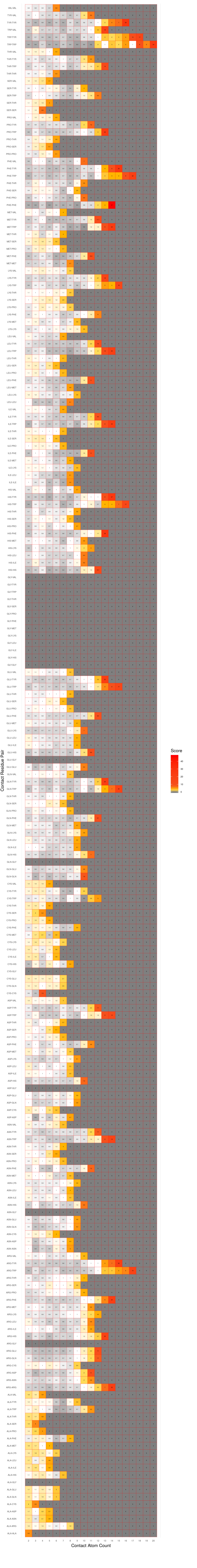

**Supplementary S6 (i).** Interacting residue pair atomic propensities at a distance threshold of 8 Å and main chain - side chain mode of interaction.

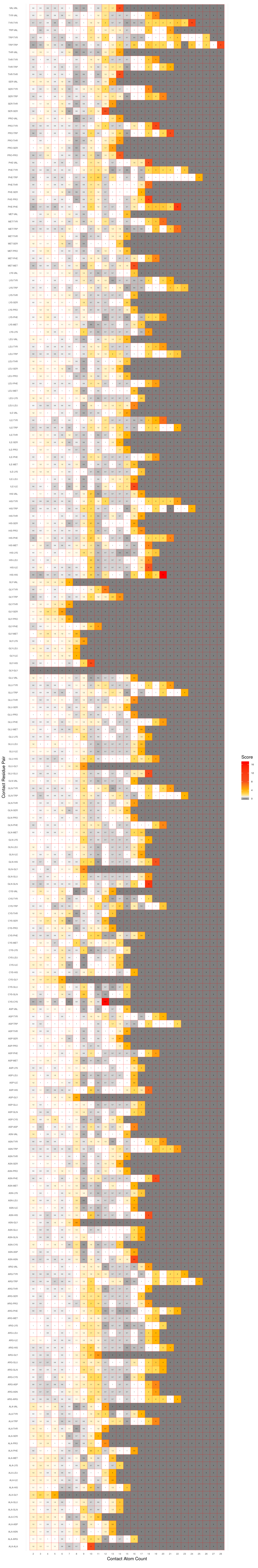

**Supplementary S7.** ROC curves for the different distance thresholds used for defining pairwise residue interactions. The Z score thresholds (operating points) along with their corresponding true positive rates and false positive rates are given in the inset table.

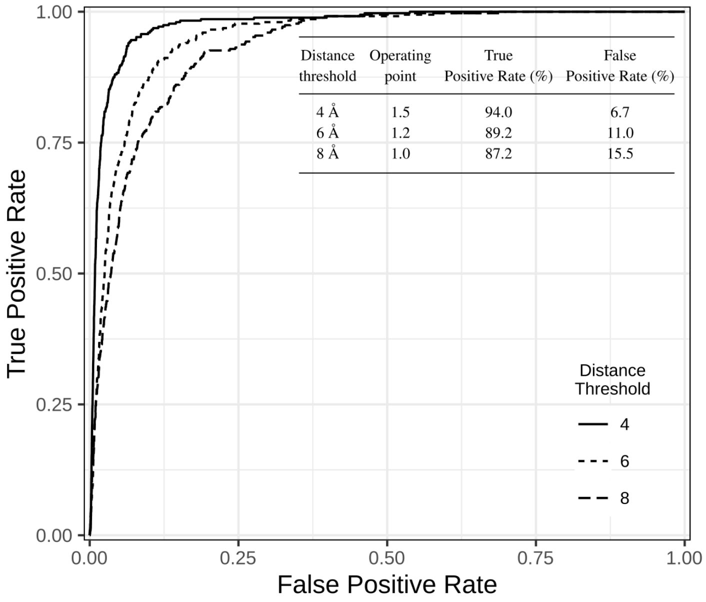

**Supplementary S8.** Benchmarking binary classification of stable associations on Dockground Docking Decoy Set 1 (Kundrotas et al., 2018) at a distance threshold of 4 Å and Z score threshold of 1.5.

| PDB code | Chain Set 1 | Chain Set 2 | Z Score |
|---|---|---|---|
| 1A2K | AB | C | 2.451 |
| 1A2Y | AB | C | 2.715 |
| 1AKJ | AB | DE | 2.386 |
| 1AVW | A | B | 1.925 |
| 1BTH | LH | P | 1.489 |
| 1BUI | A | C | 2.434 |
| 1BUI | B | C | 1.840 |
| 1BVN | P | T | 1.597 |
| 1CHO | E | I | 2.448 |
| 1DFJ | E | I | 2.202 |
| 1E96 | A | B | 2.509 |
| 1EWY | A | C | 1.788 |
| 1EZU | AB | C | 1.736 |
| 1F51 | AB | E | 1.830 |
| 1F6M | A | C | 1.947 |
| 1FM9 | A | D | 2.107 |
| 1G20 | AB | EF | 2.144 |
| 1G6V | A | K | 1.887 |
| 1GPQ | A | D | 2.311 |
| 1GPW | A | B | 2.333 |
| 1HE1 | A | C | 2.715 |
| 1HE8 | A | B | 0.889 |
| 1HXY | AB | D | 1.875 |
| 1JPS | LH | T | 2.491 |
| 1KU6 | A | B | 1.981 |
| 1L9B | LMH | C | 2.262 |
| 1MA9 | A | B | 2.339 |
| 1NBF | A | D | 2.439 |
| 1OOK | AB | G | 1.865 |
| 1OPH | A | B | 1.264 |
| 1P7Q | AB | D | 1.926 |
| 1PPF | E | I | 1.210 |
| 1R0R | E | I | 1.663 |
| 1R4M | AB | I | 2.045 |
| 1S6V | A | B | 1.250 |
| 1T6G | A | C | 2.556 |
| 1TMQ | A | B | 2.421 |
| 1TX6 | A | I | 2.044 |
| 1U7F | A | B | 2.172 |
| 1UEX | AB | C | 1.629 |
| 1UGH | E | I | 2.558 |

Continued on next page

**Table continued from previous page**

| PDB code | Chain Set 1 | Chain Set 2 | Z Score |
|---|---|---|---|
| 1W1I | A | F | 2.211 |
| 1WEJ | LH | F | 2.616 |
| 1WQ1 | R | G | 1.897 |
| 1XD3 | A | B | 2.330 |
| 1XX9 | A | CD | 1.668 |
| 1YVB | A | I | 2.138 |
| 1ZY8 | AB | K1 | 1.966 |
| 1ZY8 | AB | K2 | 2.075 |
| 2A5T | A | B | 2.577 |
| 2BKR | A | B | 2.032 |
| 2BNQ | AB | DE | 2.195 |
| 2BTF | A | P | 2.164 |
| 2CKH | A | B | 2.065 |
| 2FI4 | E | I | 1.742 |
| 2GOO | A | C | 2.414 |
| 2KAI | AB | I | 1.676 |
| 2SNI | E | I | 1.661 |
| 3FAP | A | B | 1.903 |
| 3PRO | A | C | 1.556 |
| 3SIC | E | I | 1.053 |

**Supplementary S9.** Native complex percentile ranks of Dockground Docking Decoy Set 2 (Kundrotas et al., 2018) targets evaluated using PIZSA and CIPS respectively.

| PDB code | PIZSA Percentile Rank | CIPS Percentile Rank |
|---|---|---|
| 1AY7 | 98 | 95 |
| 1BDJ | 93 | 76 |
| 1BUH | 100 | 99 |
| 1BUI | 100 | 99 |
| 1BVN | 97 | 96 |
| 1CC0 | 50 | 94 |
| 1CLV | 98 | 86 |
| 1DFJ | 100 | 100 |
| 1E6J | 97 | 99 |
| 1EER | 98 | 100 |
| 1EWY | 97 | 88 |
| 1F6M | 95 | 100 |
| 1F80 | 99 | 100 |
| 1F93 | 100 | 100 |
| 1FCC | 99 | 95 |
| 1FLE | 98 | 82 |
| 1FLT | 100 | 99 |
| 1FQ1 | 100 | 100 |
| 1FRT | 89 | 95 |
| 1G6V | 96 | 96 |
| 1G73 | 96 | 86 |
| 1GCQ | 95 | 93 |
| 1GG2 | 99 | 99 |
| 1GLB | 100 | 100 |
| 1GPQ | 99 | 83 |
| 1GPW | 100 | 100 |
| 1HE1 | 100 | 100 |
| 1HJA | 100 | 86 |
| 1HYR | 98 | 97 |
| 1I2M | 100 | 99 |
| 1I4E | 98 | 97 |
| 1ICF | 99 | 79 |
| 1IM9 | 100 | 100 |
| 1IS7 | 100 | 94 |
| 1JIW | 90 | 100 |
| 1JK9 | 100 | 89 |
| 1JTD | 100 | 100 |
| 1JZD | 100 | 100 |
| 1K5G | 100 | 100 |
| 1KGY | 95 | 48 |
| 1L9J | 69 | 78 |
| 1LB2 | 68 | 36 |
| | | Continued on next page |

**Table continued from previous page**

| PDB code | PIZSA Percentile Rank | CIPS Percentile Rank |
|---|---|---|
| 1LFD | 92 | 97 |
| 1LTX | 100 | 100 |
| 1LX5 | 99 | 98 |
| 1M63 | 100 | 71 |
| 1MG2 | 100 | 99 |
| 1ML0 | 99 | 98 |
| 1MQ8 | 100 | 89 |
| 1NBF | 100 | 100 |
| 1NQL | 80 | 100 |
| 1OC0 | 91 | 99 |
| 1OFU | 97 | 100 |
| 1PVH | 98 | 96 |
| 1QAV | 97 | 91 |
| 1R5I | 98 | 98 |
| 1R8S | 99 | 100 |
| 1RPQ | 100 | 97 |
| 1S1Q | 100 | 66 |
| 1SJH | 100 | 100 |
| 1TCO | 97 | 18 |
| 1TE1 | 99 | 96 |
| 1TMQ | 100 | 97 |
| 1U0N | 100 | 100 |
| 1UAD | 95 | 52 |
| 1UUG | 98 | 99 |
| 1V5I | 100 | 89 |
| 1V7P | 100 | 98 |
| 1VFB | 99 | 96 |
| 1VG0 | 100 | 100 |
| 1VRS | 86 | 100 |
| 1W1I | 100 | 69 |
| 1WEJ | 100 | 100 |
| 1WQ1 | 98 | 99 |
| 1WRD | 100 | 90 |
| 1X86 | 98 | 98 |
| 1XK4 | 100 | 97 |
| 1XT9 | 97 | 98 |
| 1Y8R | 100 | 14 |
| 1Y8X | 100 | 90 |
| 1YCS | 100 | 98 |
| 1YU6 | 98 | 95 |
| 1YVB | 99 | 100 |
| 1Z5Y | 99 | 100 |
| 1ZLH | 99 | 100 |
| 2A1T | 100 | 100 |
| 2A41 | 97 | 99 |

Continued on next page

**Table continued from previous page**

| PDB code | PIZSA Percentile Rank | CIPS Percentile Rank |
|---|---|---|
| 2A5D | 100 | 100 |
| 2A9K | 100 | 85 |
| 2ABZ | 92 | 100 |
| 2AQ3 | 100 | 85 |
| 2B4S | 100 | 34 |
| 2BCG | 100 | 100 |
| 2BCN | 99 | 100 |
| 2BOV | 98 | 96 |
| 2BQ1 | 97 | 100 |
| 2BWE | 96 | 99 |
| 2C0L | 100 | 96 |
| 2C2V | 98 | 100 |
| 2CH4 | 97 | 99 |
| 2D5R | 100 | 99 |
| 2EJF | 99 | 52 |
| 2G45 | 97 | 100 |
| 2GJ7 | 64 | 81 |
| 2GRX | 95 | 36 |
| 2GWF | 99 | 100 |
| 2HJ9 | 100 | 99 |
| 2HQS | 100 | 90 |
| 2HRK | 100 | 99 |
| 2I25 | 100 | 98 |
| 2IDO | 99 | 99 |
| 2IJO | 97 | 71 |
| 2IWT | 99 | 97 |
| 2J12 | 99 | 94 |
| 2NQD | 100 | 99 |
| 2NXN | 100 | 99 |
| 2O25 | 99 | 71 |
| 2O2V | 99 | 100 |
| 2O3B | 100 | 100 |
| 2O8V | 96 | 100 |
| 2OOB | 99 | 89 |
| 2PU9 | 99 | 97 |
| 2QYI | 94 | 0 |
| 2RII | 95 | 99 |
| 2SGE | 92 | 63 |
| 2UY7 | 85 | 100 |
| 2V55 | 99 | 32 |
| 2VDB | 99 | 100 |
| 2VRR | 100 | 89 |
| 2WBW | 91 | 30 |
| 2WY7 | 99 | 96 |
| 2WY8 | 99 | 99 |

Continued on next page

**Table continued from previous page**

| PDB code | PIZSA Percentile Rank | CIPS Percentile Rank |
|---|---|---|
| 2X0B | 95 | 10 |
| 2X9A | 99 | 76 |
| 2XGY | 100 | 99 |
| 2XZ1 | 100 | 100 |
| 2YVJ | 100 | 99 |
| 2ZAE | 99 | 89 |
| 2ZU0 | 99 | 100 |
| 2ZVN | 100 | 98 |
| 3A1P | 97 | 90 |
| 3A8I | 99 | 100 |
| 3AV0 | 100 | 100 |
| 3BH6 | 99 | 96 |
| 3BK3 | 100 | 100 |
| 3BP8 | 100 | 99 |
| 3BS5 | 98 | 96 |
| 3BX1 | 100 | 83 |
| 3BX7 | 100 | 99 |
| 3CBK | 100 | 94 |
| 3CII | 100 | 32 |
| 3CU1 | 95 | 7 |
| 3CVH | 100 | 80 |
| 3CW2 | 99 | 100 |
| 3D3C | 94 | 99 |
| 3D5O | 100 | 81 |
| 3D5R | 100 | 100 |
| 3DAW | 92 | 100 |
| 3E2L | 100 | 100 |
| 3F1P | 100 | 75 |
| 3F7P | 96 | 96 |
| 3FAP | 99 | 99 |
| 3FN1 | 100 | 100 |
| 3FPU | 95 | 99 |
| 3G3A | 99 | 100 |
| 3G6D | 100 | 100 |
| 3GE3 | 100 | 100 |
| 3HI6 | 100 | 98 |
| 3HZI | 100 | 90 |
| 3K1I | 100 | 93 |
| 3KLD | 98 | 99 |
| 3L1Z | 97 | 96 |
| 3L4Q | 95 | 98 |
| 3L5N | 92 | 100 |
| 3L89 | 98 | 93 |
| 3LB8 | 97 | 78 |
| 3LTF | 100 | 99 |

Continued on next page

**Table continued from previous page**

| PDB code | PIZSA Percentile Rank | CIPS Percentile Rank |
|---|---|---|
| 3LVJ | 100 | 100 |
| 3LWN | 98 | 96 |
| 3M18 | 100 | 66 |
| 3MJ7 | 99 | 100 |
| 3MZW | 100 | 99 |
| 3OED | 99 | 90 |
| 3OJ4 | 99 | 92 |
| 3ONA | 92 | 95 |
| 3ONL | 97 | 100 |
| 3OUN | 99 | 79 |
| 3PNL | 100 | 88 |
| 3PRO | 100 | 74 |
| 3PRP | 100 | 100 |
| 3PV6 | 98 | 88 |
| 3QC8 | 100 | 100 |
| 3QLU | 100 | 92 |
| 3R66 | 100 | 100 |
| 3R9A | 100 | 99 |
| 3RJ3 | 100 | 57 |
| 3RNK | 99 | 97 |
| 3RRM | 100 | 90 |
| 3S36 | 100 | 99 |
| 3S3X | 97 | 100 |
| 3S5L | 98 | 100 |
| 3T1Q | 100 | 100 |
| 3T5G | 97 | 27 |
| 3TG1 | 99 | 100 |
| 3UAI | 100 | 100 |
| 3ULQ | 100 | 69 |
| 3VLB | 100 | 98 |
| 3VXU | 82 | 95 |
| 3W31 | 99 | 100 |
| 3WKT | 98 | 100 |
| 3WWK | 100 | 100 |
| 3WWN | 100 | 96 |
| 3ZN6 | 100 | 92 |
| 3ZNZ | 99 | 94 |
| 4BI8 | 100 | 100 |
| 4BMP | 100 | 98 |
| 4BOS | 100 | 100 |
| 4BOZ | 99 | 100 |
| 4BWS | 94 | 94 |
| 4C0O | 100 | 98 |
| 4C6T | 99 | 93 |
| 4CPA | 93 | 92 |

Continued on next page

**Table continued from previous page**

| PDB code | PIZSA Percentile Rank | CIPS Percentile Rank |
|---|---|---|
| 4CT4 | 100 | 100 |
| 4CU4 | 100 | 41 |
| 4DI3 | 100 | 99 |
| 4DN4 | 97 | 100 |
| 4DSS | 100 | 100 |
| 4E2I | 100 | 100 |
| 4EJX | 100 | 61 |
| 4EMJ | 99 | 99 |
| 4ETQ | 100 | 100 |
| 4ETW | 99 | 100 |
| 4EXT | 100 | 89 |
| 4F7G | 100 | 88 |
| 4FA8 | 100 | 89 |
| 4FI3 | 100 | 92 |
| 4FME | 95 | 89 |
| 4GED | 100 | 98 |
| 4GMJ | 100 | 97 |
| 4GOJ | 99 | 100 |
| 4HCN | 99 | 87 |
| 4HH3 | 100 | 90 |
| 4HMY | 95 | 98 |
| 4HQP | 99 | 100 |
| 4HRE | 100 | 100 |
| 4HX3 | 100 | 60 |
| 4I5L | 100 | 100 |
| 4ICG | 99 | 91 |
| 4IHF | 100 | 100 |
| 4ILH | 100 | 100 |
| 4J38 | 100 | 87 |
| 4J4L | 99 | 100 |
| 4JAV | 100 | 99 |
| 4JCV | 96 | 98 |
| 4JHP | 99 | 96 |
| 4JO9 | 100 | 96 |
| 4JQW | 97 | 70 |
| 4JX1 | 100 | 100 |
| 4K0V | 80 | 96 |
| 4K71 | 100 | 100 |
| 4KEH | 100 | 100 |
| 4KI1 | 100 | 100 |
| 4KR0 | 100 | 100 |
| 4KXZ | 100 | 96 |
| 4KYI | 99 | 100 |
| 4L41 | 100 | 100 |
| 4LHU | 100 | 100 |

Continued on next page

**Table continued from previous page**

| PDB code | PIZSA Percentile Rank | CIPS Percentile Rank |
|---|---|---|
| 4LNU | 100 | 100 |
| 4LRZ | 100 | 88 |
| 4LSX | 100 | 100 |
| 4LW4 | 98 | 99 |
| 4LX0 | 98 | 73 |
| 4M3K | 100 | 93 |
| 4M7L | 91 | 96 |
| 4MDK | 93 | 90 |
| 4MNE | 100 | 100 |
| 4MRT | 99 | 100 |
| 4N6O | 96 | 72 |
| 4NBE | 100 | 100 |
| 4NIF | 100 | 97 |
| 4NYI | 100 | 91 |
| 4NZL | 91 | 91 |
| 4O4B | 96 | 100 |
| 4P2A | 99 | 99 |
| 4P4H | 100 | 84 |
| 4P4Q | 100 | 100 |
| 4P69 | 100 | 98 |
| 4PDC | 100 | 92 |
| 4PJ2 | 96 | 98 |
| 4PJ8 | 100 | 99 |
| 4POU | 99 | 97 |
| 4PW9 | 100 | 60 |
| 4QD2 | 100 | 93 |
| 4QTI | 98 | 100 |
| 4R62 | 85 | 1 |
| 4R9Y | 100 | 100 |
| 4RF0 | 100 | 81 |
| 4RIX | 96 | 90 |
| 4RWS | 100 | 48 |
| 4S10 | 99 | 99 |
| 4TXV | 100 | 78 |
| 4V2C | 74 | 32 |
| 4WLR | 100 | 99 |
| 4WM0 | 93 | 94 |
| 4WZA | 100 | 100 |
| 4XHU | 100 | 100 |
| 4XIG | 100 | 42 |
| 4XR8 | 100 | 91 |
| 4XS0 | 100 | 92 |
| 4YEB | 100 | 97 |
| 4ZFR | 100 | 95 |
| 5AUP | 100 | 96 |

Continued on next page

**Table continued from previous page**

| PDB code | PIZSA Percentile Rank | CIPS Percentile Rank |
|---|---|---|
| 5BMU | 100 | 100 |
| 5BVP | 100 | 100 |
| 5C3I | 100 | 98 |
| 5C7X | 100 | 95 |
| 5CEC | 100 | 93 |
| 5CRA | 99 | 98 |
| 5CVO | 100 | 89 |
| 5E0K | 100 | 99 |
| 5E8D | 100 | 56 |
| 5EG3 | 91 | 100 |

**Supplementary S10 (a).** Identification of near native decoys from the CAPRI Score_set. Percentage of near-native decoys (y-axis) ranked in the top 1%, 10%, 20%, 30%, 40% and 50% (x-axis) of all decoys in different CAPRI targets. Lines are coloured according to the near native decoy category: high (green), medium (blue) or acceptable (pink). Counts are provided for every data point.

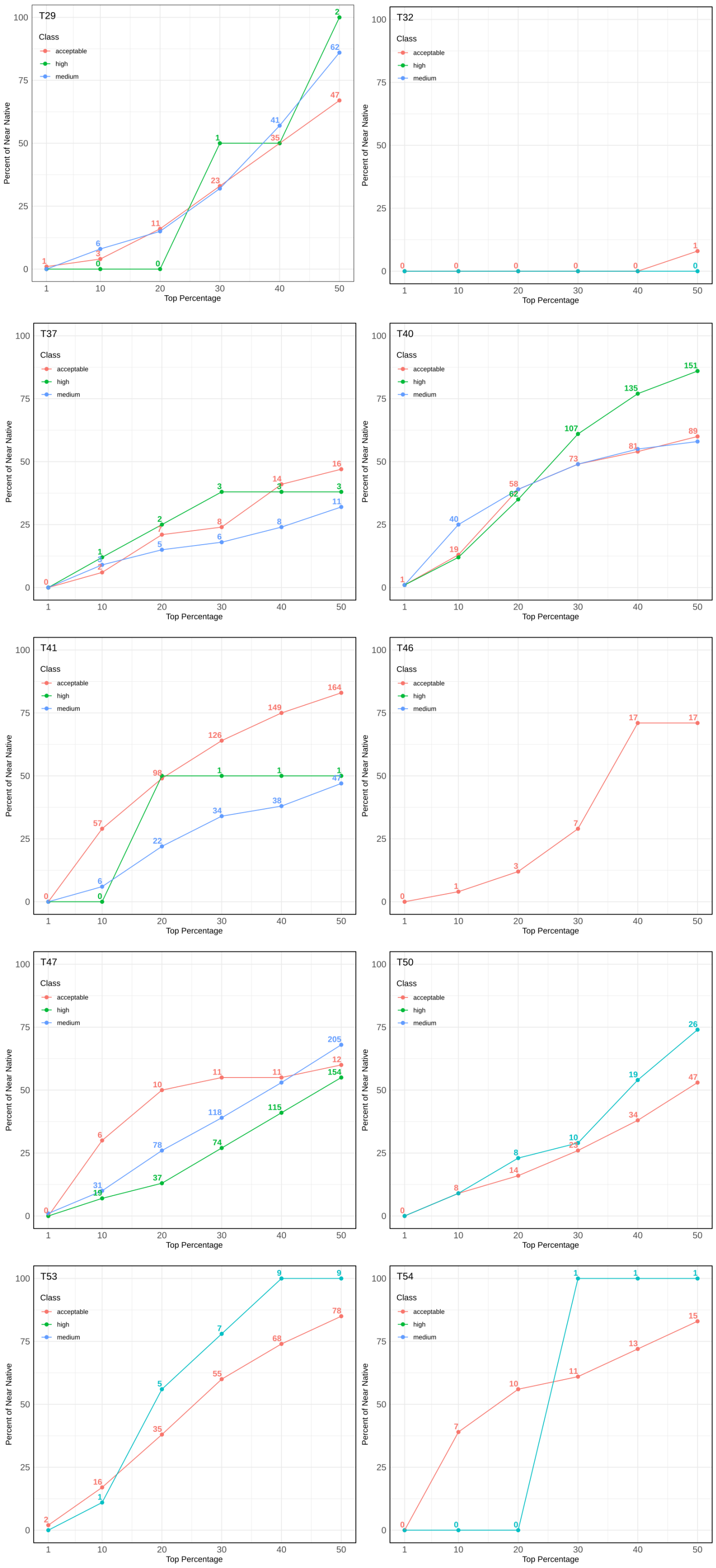

**Supplementary S10 (b).** List of top ranked high, medium and acceptable near native decoys from the CAPRI Score_set. The number and percentage of near native decoys from each category ranked in the top 1%, 10%, 20%, 30%, 40% and 50% of decoys as scored with PIZSA and CIPS are reported.

| Quality | Top % | Number of near natives | | Percent of near natives | |
|---|---|---|---|---|---|
| | | PIZSA | CIPS | PIZSA | CIPS |
| | 1 | 2 | 0 | 0.4 | 0.0 |
| | 10 | 41 | 20 | 8.8 | 4.3 |
| **High** | 20 | 102 | 97 | 21.9 | 20.8 |
| | 30 | 186 | 191 | 39.9 | 41.0 |
| | 40 | 255 | 297 | 54.7 | 63.7 |
| | 50 | 311 | 361 | 66.7 | 77.5 |
| | 1 | 4 | 10 | 0.6 | 1.4 |
| | 10 | 90 | 111 | 12.5 | 15.4 |
| **Medium** | 20 | 193 | 243 | 26.8 | 33.8 |
| | 30 | 279 | 364 | 38.8 | 50.6 |
| | 40 | 367 | 462 | 51.0 | 64.2 |
| | 50 | 458 | 548 | 63.6 | 76.1 |
| | 1 | 4 | 18 | 0.6 | 2.5 |
| | 10 | 119 | 93 | 16.7 | 13.1 |
| **Acceptable** | 20 | 246 | 177 | 34.6 | 24.9 |
| | 30 | 337 | 306 | 47.4 | 43.0 |
| | 40 | 422 | 410 | 59.4 | 57.7 |
| | 50 | 488 | 519 | 68.6 | 73.0 |

**Supplementary S10 (c).** Classification of near native decoys as stable associations. Classification performance was tested on the CAPRI Score_set decoys with 1,896 near native complexes (positives; high, medium and acceptable decoys) and 16,676 decoy complexes (negatives). Near native complexes classified as stable or unstable associations are identified as true positives (TP, near native complexes with Z Score $> 1.5$) or false negatives (FN, near native complexes with Z Score $<= 1.5$) respectively. Decoy complexes classified as stable or unstable associations are identified as false positives (FP, decoy complexes with Z Score $> 1.5$) or true negatives (TN, decoy complexes with Z Score $<= 1.5$) respectively. PIZSA classified the complexes with a True Positive Rate (TPR) of 0.63, False Negative Rate (FNR) of 0.37, True Negative Rate (TNR) of 0.62, False Positive Rate (FPR) of 0.34 and Accuracy of 0.62.

| | Positive | Negative |
|---|---|---|
| **True** | 1,201 | 10,363 |
| **False** | 6,313 | 695 |
